# Repeated global adaptation across plant species

**DOI:** 10.1101/2024.04.02.587814

**Authors:** Gabriele Nocchi, James R. Whiting, Samuel Yeaman

## Abstract

Global adaptation occurs when all populations of a species undergo selection toward a common optimum. This can occur by a hard selective sweep with the emergence of a new globally advantageous allele that spreads throughout a species’ natural range until reaching fixation. This evolutionary process leaves a temporary trace in the region affected, which is detectable using population genomic methods. While selective sweeps have been identified in many species, there have been few comparative and systematic studies of the genes involved in global adaptation. Building upon recent findings showing repeated genetic basis of local adaptation across independent populations and species, we asked whether certain genes play a more significant role in driving global adaptation across plant species. To address this question, we scanned the genomes of 17 plant species to identify signals of repeated global selective sweeps. Despite the substantial evolutionary distance between the species analysed, we identified several gene families with strong evidence of repeated positive selection. These gene families tend to be enriched for reduced pleiotropy, consistent with predictions from Fisher’s evolutionary model and the cost of complexity hypothesis. We also found that genes with repeated sweeps exhibit elevated levels of gene duplication. Our findings contrast with recent observations of increased pleiotropy in genes driving local adaptation, consistent with predictions based on the theory of migration-selection balance.

**Significance:** Global adaptation occurs when a species undergoes selection toward a common optimum throughout its natural range. While instances of global adaptation are widespread in the literature, there is a shortage of comparative studies aimed at understanding its genetic architecture and how it contrasts with that of local adaptation. This research compares global selective sweeps across 17 plant species to uncover the attributes of the genetic loci repeatedly involved in adaptation. We show that global adaptation tends to rely on genes with reduced pleiotropy and is characterized by increased levels of gene duplication. This finding contrasts with recent observations of increased pleiotropy in genes driving local adaptation, reflecting the opposing dynamics underlying these two evolutionary processes.

## Introduction

Plant species occupy a wide range of niches, adopt very different life history strategies, and inhabit environments with drastically different biotic pressures (1). Due to their sessile nature, plants must contend with the biotic and abiotic stresses they encounter directly in their environment, therefore phenotypic plasticity and genomic adaptation are of critical importance in the plant kingdom (1). To no surprise, local adaptation has been more commonly detected in plants than in animals (2,3,4,5).

Local adaptation occurs when a species inhabits a heterogeneous environment with spatial variability in the optimal phenotype. This can lead to the evolution of spatially differentiated genotypes along environmental gradients that exhibit fitness trade-offs when transplanted, with local genotypes providing higher fitness than foreign (4,5,6). By contrast, global adaptation occurs when all populations of a species experience selection towards the same optimum, resulting in the gradual refinement of a trait that is advantageous throughout all the environments inhabited by a species, such as the evolution of opposable thumbs in ancestral humans (6). While both global and local adaptation involve positive selection, the tension between migration and divergent natural selection inherent in local adaptation can result in different predictions about the evolution of genetic architecture. Local adaptation will tend to involve alleles with larger effects and more tightly linked than global adaptation, as such architectures are better able to resist the homogenizing effect of migration (6). However, while this is a clear theoretical prediction, it has not been extensively tested using empirical data.

At the molecular level, global adaptation can occur via hard or soft selective sweeps or through more subtle shifts in allele frequency at many loci (7). With a hard selective sweep (8,9,10) a new globally advantageous mutation rapidly spreads throughout a species’ natural range until it reaches fixation. During this process, the affected genomic region displays a distinctive signature marked by diminished genetic diversity and increased homozygosity, a shift of the site frequency spectrum (SFS) toward low and high-frequency variants (11,12), and particular patterns of linkage disequilibrium (LD) characterized by elevated LD on each side of the selected site and decreased LD between sites located on different sides (13). This trace can be detected by scanning intraspecific genetic data, and various population genomics methods have been developed to detect sweeps (10,14,15). After fixation of the beneficial variant, new mutations and recombination in the region slowly decay the genomic signature typical of a hard selective sweep, therefore genomic scans can only detect hard sweeps within a restricted time frame (14). Global adaptation can also arise through positive selection on standing genetic variation, which tends to result in soft selective sweeps (16) or more subtle allele frequency shifts at many loci (7). When a beneficial variant is already segregating in a population before being subjected to positive selection, the selective process does not affect linked neutral polymorphisms as much as in hard-selective sweeps, making the detection of soft-sweeps much more challenging (10,16).

While selective sweeps have been reported in many species across both plants and animal kingdoms (17,18,19,20,21,22,23,24,25,26,27,28,29,30,31), there has been limited comparative and systematic genome-wide study of the repeatability of gene involvement in global adaptation across multiple species. On the other hand, several studies have demonstrated that the same genetic loci are observed repeatedly driving local adaptation in different populations or species (32,33,34,35,36,37,38,39,40,41). Whether adaptation is local or global, one important factor affecting the extent of repeatability is genotypic redundancy. If a given adaptive trait is characterized by limited genotypic redundancy (6), few loci are available to produce the mutations needed to reach the phenotypic optimum, therefore more repeatability is expected. In such cases, adaptive loci usually have large effects, a pattern that has recently emerged for several cases of local adaptation (42). On the other hand, if a trait is highly polygenic and driven by numerous alleles of small and interchangeable effect (6,43), a multitude of genotypes could potentially yield the same optimum phenotype. In such cases, fewer loci are expected to exhibit repeated contributions to adaptation in independent bouts of evolution across species or lineages.

As repeated genetic basis of adaptation has been identified among different populations and in closely related species, it is intriguing to assess whether repeatability is observed at greater phylogenetic distances (41, 44). It is expected that more recently diverged lineages will demonstrate a higher degree of shared gene utilization in adaptation, owing to limited functional differentiation, increased allele sharing, comparable genomic architecture, and similar life histories/adaptive strategies (41). Also, while factors such as pleiotropy, mutability, and average mutation effect size likely vary among genes and would be theoretically predicted to affect the repeatability of adaptation, there has been little systematic study of the importance of such factors in empirical datasets.

To address these questions, we retrieved publicly available high-quality whole-genome sequencing (WGS) data from thousands of individuals from 17 natural plant and forest trees species distributed across four continents, including woody and herbaceous angiosperms species (Fig. 1 and SI Appendix, Table S1). We employed linkage disequilibrium-based genomic scans for selective sweeps (45) within each species to identify putative genes under positive selection, and then used *PicMin* (46) to identify genes that are enriched for strong sweep signatures across multiple species. We explored the attributes of genes with repeated global sweeps by examining the pleiotropic potential and gene duplication levels of the repeatedly swept gene families identified. Our assessment of pleiotropy leveraged available gene expression data from *Arabidopsis thaliana* and *Medicago truncatula* to generate different pleiotropy measures based on gene tissue specificity and position within gene co-expression networks. We also assessed levels of gene duplication based on our orthology assignment. We contextualized these findings in view of Fisher’s model of evolution and the cost of complexity hypothesis (47, 48), as well as recent theoretical work on migration-selection balance (6). To contrast our results for the architecture of global adaptation with that of local adaptation, we compare our observations to results from a related study on local adaptation (49), which employed the same bioinformatic methods and incorporated 14 of the 17 datasets used in our analysis.

**Fig. 1.**
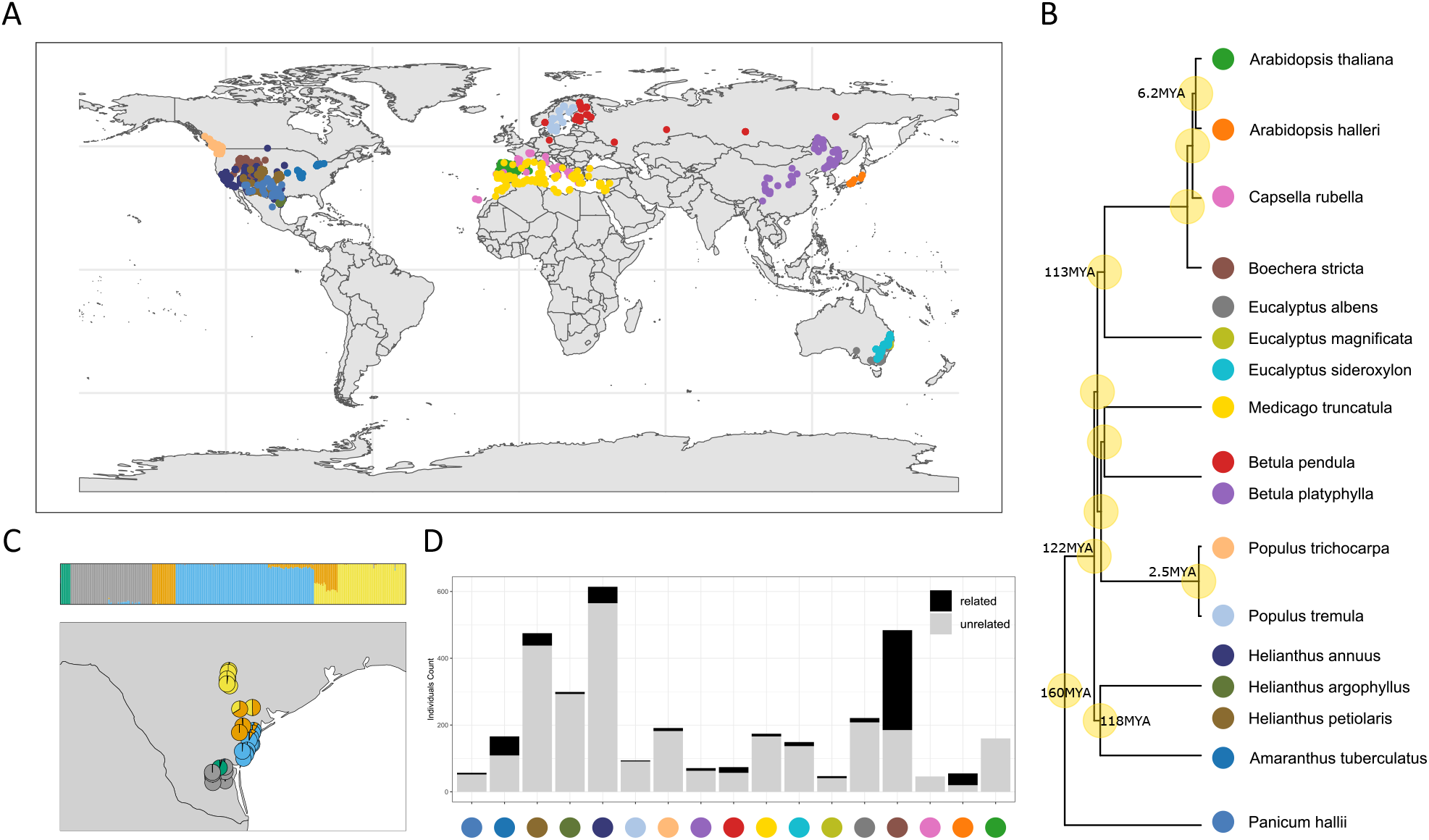
Geographical distribution and relatedness within and among the study species. (A) Sampling locations of the 17 datasets included in the study. (B) Time-calibrated phylogenetic tree (retrieved from https://timetree.org/) of the 17 datasets, based on 12 reference species. (C) *fastSTRUCTURE* ancestry pie plot (K = 5) of the *Helianthus argophyllus* dataset in Texas (USA), showing substantial sub structuring. (D) Relatedness filtering summary bar plot by dataset. Datasets labelled by corresponding colour from B.

## Results

### Datasets assembly and filtering

Whole-genome sequencing data (WGS) from 17 non-domesticated and non-invasive angiosperm species, including both herbaceous plants and forest trees, was retrieved from SRA/ENA (Fig. 1 and SI Appendix, Table S1). These datasets included exclusively natural populations and ranged from closely related sister species known to hybridize (50) to species separated by up to 160 million years of evolution (Fig. 1B).

Single nucleotide polymorphisms (SNPs) were identified by applying a uniform pipeline to all datasets for consistency, with filtering steps to exclude singleton SNPs and individuals with high relatedness (SI Appendix, Fig. S1; see *Methods* for details). The SNP calling and filtering process resulted in between 20 and 565 individuals per dataset, and between 667,641 and 50,268,965 SNPs (Fig. 1D and SI Appendix, Table S1). We assessed population structure and found a wide range of patterns, with some species showing nearly panmictic structure across wide spatial scales (e.g. *Eucalyptus albens*; SI Appendix, Fig. S2) and others exhibiting substantial sub structuring, even across small spatial scales (e.g. *Helianthus argophyllus*) (Fig. 1C and SI Appendix, Fig. S2). In most species, the range of samples included in our study is sufficient to either cover multiple populations with detected genetic structure (SI Appendix, Fig. S2) or span a substantial portion of the overall species range (>20%), as estimated using data from GBIF (SI Appendix, Fig. S3). The only exception is *P. tremula*, which exhibited a panmictic genetic structure and had samples spanning approximately 11.3% of its range (SI Appendix, Fig. S2 and Fig. S3). Despite this, it included several sampling sites scattered over approximately 1,100 km of latitude comprising a large number of individuals (SI Appendix, Fig. S2 and Table S1), so it still should be seen as representative of capturing patterns of adaptation across a broad range of space. It is important to note that the native range estimation of the species was likely overestimated due to the possible presence of erroneous observations, observations of planted specimens and observations within invasive ranges in the GBIF data.

### Orthology inference

The phylogenetic relationship among the genes of the 12 species with reference genome assemblies was inferred with *Orthofinder2* (51), which was used to group genes into orthology groups, or orthogroups. These included both orthologous genes, which are homologous genes separated by a speciation event, and paralogous genes, which are homologous genes diverged from a within-species duplication event. In total, *Orthofinder* assigned 376,881 genes out of 415,552 total genes to 31,582 orthogroups. These included 9,521 orthogroups specific to individual species, 7,919 orthogroups with representation from all species, and 633 single-copy orthogroups. Subsequently, this assignment was refined by excluding cases where a given species had more than 10 paralogues within a single orthogroup (but retaining the orthogroup in other species with fewer than 10 paralogues). We also excluded any gene with insufficient sequencing coverage within a given species, and excluded any orthogroups with representation in fewer than 7 species. These filtering steps resulted in a final set of 13,268 orthogroups for subsequent analysis, which exhibited low levels of paralogy and high species inclusion (Fig. 2). The mean and median number of paralogues per orthogroup was 1.72 and 1.67 per species respectively, while mean and median species number per orthogroup was 15.37 and 15.42 (per species). Notably, *B. stricta* yielded results for significantly fewer orthogroups compared to the other species, owing to insufficient sequencing coverage of the data (52) across many genic regions (Fig. 2 and SI Appendix,Table S1). Overall, the topology of the tree inferred with *Orthofinder2* largely matched the species tree derived from *TimeTree* (https://timetree.org/), with the exception of *M. truncatula* and *E. grandis. Orthofinder2* placed *M. truncatula* closer to the Brassicaceae family, relative to *E. grandis* (Fig. 2D and SI Appendix, Fig. S4). It is worth noting that *TimeTree* estimates divergence between species using a simple average across published time estimates from scientific literature (53), therefore a minor mismatch with the protein sequences-based tree inferred with *Orthofinder2* is not a reason of concern.

**Fig. 2.**
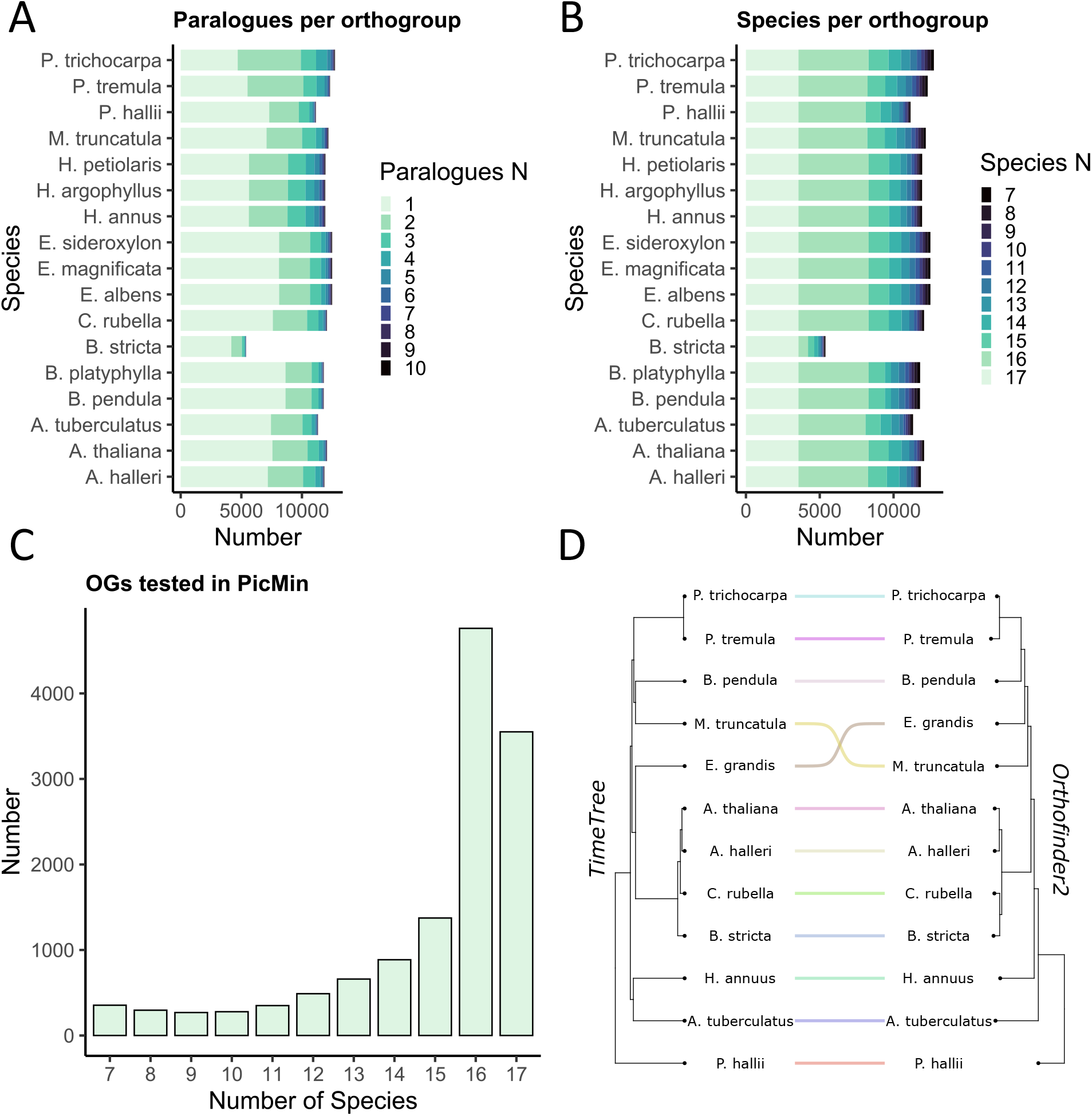
Orthology assignment summary of the final set of 13,268 orthogroups. (A) Bar plot of the number of paralogues per orthogroup for each species. (B) Bar plot of the number of species included in each orthogroup, for the orthogroups of each species. (C) Distribution of the 13,268 tested orthogroups across species number. (D) Comparison between the *TimeTree* and *Orthofinder2* phylogenies, each based on 12 reference species.

### Repeated selective sweeps

We used *OmegaPlus* (45) to scan for global selective sweeps within species followed by *PicMin* (46) to identify orthogroups with repeated sweep signatures across several species. After performing a False Discovery Rate (FDR) correction to the p-values from *PicMin* based on the number of orthogroups tested, we detected 33 orthogroups with significant signatures of repeated sweeps, at q < 0.1 (Fig. 3A).

**Fig. 3.**
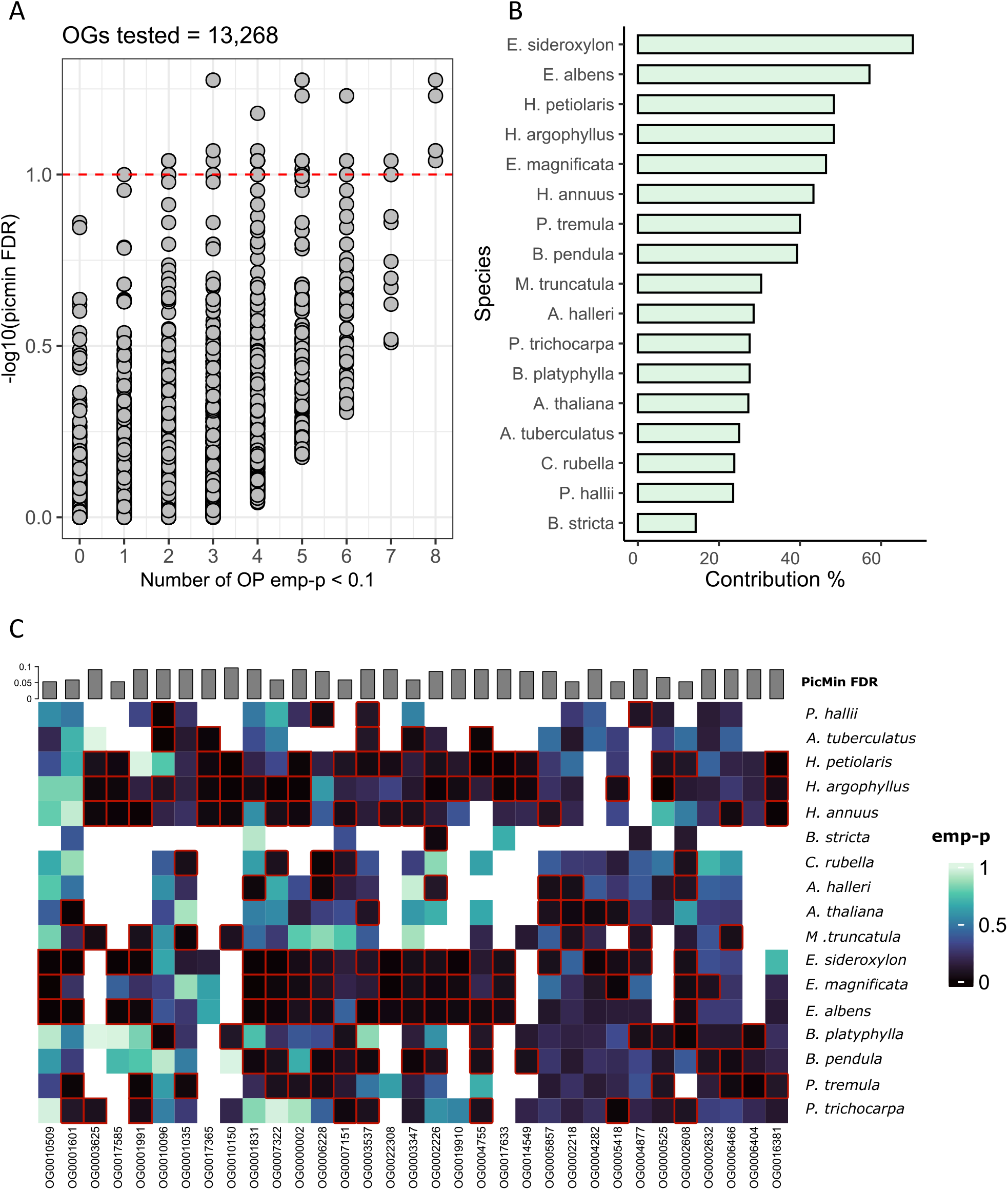
Evidence for repeated selective sweeps across multiple species. (A) Distribution of *PicMin* FDR (-log10 *PicMin* FDR on the Y axis) for the 13,268 tested orthogroups, ordered by number of putative driving genes on the X axis (number of *OmegaPlus* emp-p < 0.1). Points above the red line have FDR < 0.1. (B) Species contribution to *PicMin* top candidates (FDR < 0.1), calculated for each species as: [number of empirical p-values < 0.1 in the *PicMin* FDR < 0.1 orthogroups]/[total number of orthogroups with *PicMin* FDR < 0.1 tested]. (C) Heatmap of *OmegaPlus* empirical p-values for the 33 candidate orthogroups (*PicMin* FDR < 0.1); driving genes cells (emp-p < 0.1 – black cells) are outlined in red, species are ordered by phylogenetic distance along the Y axis. A white cell indicates that an orthogroup was not tested in a species.

Our application of *PicMin* tests for an enrichment of strong sweep signatures across multiple species but does not perform a test of which species contribute to the signal. To explore any patterns in the species driving these results, we classify any gene with an empirical p-value of < 0.1 as a “driving gene”. Contribution towards the signatures of repeatability in the 33 OGs varied between species, ranging from approximately 14% (*B. stricta*) driving genes to about 68% (*E. sideroxylon*) (Fig. 3B). The repeatability signal is scattered throughout the phylogeny, with low *OmegaPlus* empirical p-values present in every species tested (Fig. 3C). However, two clusters of closely related species (*Eucalyptus spp*. and *Helianthus spp*.) were top contributors based on the count of driving genes and displayed a denser heat signature indicating an enrichment of driving genes (Fig. 3B, Fig. 3C and SI Appendix, Fig. S5). Despite this, results for only a single orthogroup appeared to be driven solely by species from the *Eucalyptus* genus, OG0010509, and none was solely driven by the *Helianthus* genus (Fig. 3C and SI Appendix, Fig. S5). However, in four orthogroups (OG0017633, OG0019910, OG0022308, OG0017585), driving genes were found exclusively in multiple *Eucalyptus* species in conjunction with multiple *Helianthus* species (Fig. 3C and SI Appendix, Fig. S5).

It is biologically expected that closely related species might evolve similarly and therefore might be more likely to experience sweeps in the same genes or genes families, particularly if they occupy an overlapping habitat and experience the same, or similar, selective pressures (41, 54). Alternatively, sweeps detected on the same gene of closely related species of the same genus could have originated from a single sweep that was then introgressed across these hybridizing species. However, it is also possible that such signatures are bioinformatic artefacts, as a single reference genome was used to call variants in the species within each of the *Helianthus* and *Eucalyptus* genera, which means that any assembly errors could yield anomalous LD signatures that might confound *OmegaPlus* in each of the species. Consequently, the same errors could systematically propagate across closely related species sharing the same reference and potentially drive some of the observed *PicMin* repeatability.

To test whether this was a source of error, we assessed whether phylogenetic distance correlated with the amount of overlap in the driving genes between pairs of species, which we calculated as the ratio of observed versus expected overlap, according to the expectation of a hypergeometric distribution, and found no significant correlation (*Pearson’s r* = 0.05, *p-value* = 0.57; *Spearman’s rho* = -0.05, *p-value* = 0.6) (SI Appendix, Fig. S6A). Overall, the observed overlap in driving genes between closely related species did not appear to deviate from the expectation differently than that between more distantly related species. Further assessment of the average phylogenetic distance between driving genes species showed that all the candidates, except one (OG0010509), are characterized by considerable mean phylogenetic distances between the species with driving genes (SI Appendix, Fig. S6B). This confirms that despite an apparent enrichment of low empirical p-values in *Helianthus* and *Eucalyptus* species, the overall repeatability signal is driven by genes spread throughout the analysed phylogeny.

To preclude spurious contributions from closely related species arising from bioinformatic artefacts, we conducted follow-up repeatability testing by running *PicMin* with only one species from each of these two genera and taking the union of significant hits (FDR < 0.5) across runs, evaluating all nine of the 13-species combinations [1 *Helianthus* + 1 *Eucalyptus* + all other species]. While this restricted analysis cannot detect true signals of repeated sweeps within a genus, it does not suffer from the risk of being driven by bioinformatic errors propagating due to closely related species sharing the same reference genome. Despite reduced power due the overall lower number of species included, this follow-up analysis successfully retrieved 15 of the 33 originally identified candidate orthogroups (FDR < 0.1) (SI Appendix, Table S2). Finally, we tested the robustness of *PicMin* against statistical artifacts and potential biases introduced by factors such as gene length, variations in *OmegaPlus* settings and scanning strategies, shared recombination landscape and background selection (please see SI Appendix, Supporting Text – Possible biases and artifacts).

Next, we explored the functional characteristics of the highest confidence set of 15 orthogroups with repeated sweep signatures, using the annotations of *A. thaliana* (55) and *M. truncatula* (56) genes (SI Appendix, Table S2). These orthogroups included several genes with pivotal roles in biotic and abiotic stress responses. For instance, TPS14 (TERPENE SYNTHASE 14) is a key protein involved in terpene metabolism. Terpenes are metabolites involved in plant defence against both pathogens and herbivores, and are responsible for the attraction of beneficial organisms such as pollinators and seed dispersers (57, 58). Similarly, TGSL12 (CALLOSE SYNTHASE 3) is a protein that participates in callose metabolism, a polysaccharide synthesized in the cell wall of a variety of higher plants in response to pathogens infections and abiotic stress (59). The extensively studied cytochrome P450, belongs to a large family of proteins involved in NADPH- and O_2_-dependent hydroxylation reactions across all domains of life and is closely linked to environmental stress response in plants (60). Notably ATERF019 (ERF019, ERF19), is a critical transcription factor involved in the response to a range of stressors, such as drought, osmotic, and oxidative stress, as well as bacterial and fungal infection (61, 62). Intriguingly, this gene has been found to negatively regulate plant resistance to *Phytophthora parasitica*, underscoring its significance in mitigating the impact of destructive plant pathogens causing significant crop losses worldwide (61). Another notable protein group identified was the Plant U-box type E3 ubiquitin ligase (PUB62, PUBs), which encompasses proteins with diverse functions in abiotic and biotic stress responses and is also associated with pollen self-incompatibility (63).

In addition to stress responses, several orthogroups with signatures of repeated sweeps play roles in plant growth and development. For instance, the clathrin adaptor complexes medium subunit family protein is involved in clathrin-mediated endocytosis (CME). This process regulates various aspects of plant development, such as hormone signaling, and has also been linked to response against environmental stresses (64). Interestingly, the machinery of CME has been shown to be evolutionarily conserved in plants (64). Furthermore, we identified the p300/CBP acetyltransferase-related protein, which is linked to cell differentiation, growth, and homeostasis. Notably, this protein has a 600 amino acid C-terminal segment which appears highly conserved across plants and animals, suggesting an essential role in multicellular organisms (65). Other key players in growth and development included ATSPLA2-ALPHA (PHOSPHOLIPASE A2-ALPHA), which is involved in various growth-related processes (66), and ROXY1/ROXY2 of the CC-type glutaredoxin (ROXY) family, which is a group of proteins crucial for organogenesis, particularly anther development (67). Lastly, among our top candidates was ATIPT4 (ARABIDOPSIS THALIANA ISOPENTENYLTRANSFERASE 4), a protein that plays a significant role in cytokinin biosynthesis. Cytokinins are hormones essential for regulating various aspects of plant growth and development (68).

### Spatial scale of adaptation

Selective sweeps are typically interpreted as evidence of a mutation spreading throughout the range of an entire species (i.e. global adaptation), but they can also occur within a restricted portion of the range due to local adaptation. It is possible that some of the signatures of repeated sweeps we observed could be driven by strong local adaptation within a subsection of the species range, rather than global adaptation. To explore this possibility, we estimated *F*_ST_ for each SNP within each species and took the average across all SNPs within each gene. Driving genes in orthogroups with repeated sweep signatures displayed low to average *F*_ST_ within species, with few exceptions, suggesting that the signatures of repeated selective sweeps were driven by global, rather than local adaptation (SI Appendix, Fig. S7 and Fig. S8). Even though we observed few driving genes with high *F*_ST_ (8 genes out of 78 fell within the top 10% of *F*_ST_ values genome wide; SI Appendix, Fig. S8), it is worth noting that a mutation making a global selective sweep can generate transient genetic differentiation as *F*_ST_ outliers, so even the relatively rare cases of high *F*_ST_ that we observed within species could still have been driven by global adaptation (69).

### Genes with repeated sweeps have low pleiotropy

We explored the pleiotropic characteristics of orthogroups with repeated sweep signatures to assess the genetic architecture of global adaptation in comparison to the classical theoretical expectation based on Fisher’s geometric model of universal pleiotropy (47,48), other empirical studies (32,33,34,35,36,37,38,39,70,71,72,73,74), as well as more recent findings on the architecture of local adaptation derived from a large comparative study across plant species sharing the same methods and majority of datasets, including the same SNPs sets (49).

To evaluate the amount of pleiotropy in genes with repeated selective sweeps, we utilized gene expression data for *A. thaliana* and *M. truncatula* genes sourced from Expression Atlas (75) and ATTED-II (76). We estimated two kinds of gene property related to pleiotropy: A) tissue specificity of gene expression (77) and B) statistics describing a gene’s position/importance within a gene co-expression network (78). For tissue specificity (A), we employed the τ metric (79) of *A. thaliana* genes, which should be inversely proportional to pleiotropy; highly tissue-specific genes are likely to influence fewer processes compared to genes expressed across several tissues. Tissue specificity has been found to be positively associated with rates of molecular evolution (80). Using gene co-expression networks (B), we estimated pleiotropy based on four centrality measures (see *Methods* for details): degree, strength, closeness, and betweenness (81). Node degree signifies the number of nodes connected to a gene and thus the number of co-expressed genes; node strength represents the combined weights of these connections; closeness reflects a node’s capacity to interact with all other nodes, even those not directly connected to it, and hence its co-expression capability across the entire network; betweenness indicates a node’s ability to act as a bridging node in the network, linking co-expression subnetworks together. Each of these measures should be positively correlated to pleiotropy (82,83,84). Genes with high centrality measures in co-expression networks are hubs; hence, their expression affects many other genes and processes, potentially imposing an evolutionary constraint as changes in them can more likely be lethal or have strong effects. Conversely, genes with low centrality measures can be considered less pleiotropic, with changes in them affecting fewer processes (82,83,84).

To evaluate the amount of pleiotropy in the candidate orthogroups with repeated sweep signatures, each of the above measures was calculated for each orthogroup based on the metrics described above, using data from *A. thaliana* and *M. truncatula* (see *Methods* for details). We assessed pleiotropy in the candidate set including 33 Orthogroups (FDR < 0.1), as well as within a larger set of 107 orthogroups with a more relaxed false discovery rate cutoff (FDR < 0.2). While using a more relaxed FDR results in more false positives, it also allows a larger number of true positives, which improves the power to test patterns in pleiotropy for genes involved in global adaptation. Consistent with the Fisher-Orr model of evolution (47,48), both sets of results showed the same pattern: pleiotropy appeared significantly lower in the candidate orthogroups compared to random expectation (Fig. 4A, Fig. 4B). All measures used as proxies for pleiotropy, except *Medicago* betweenness, showed either a significant (*p* < 0.05) association with decreased pleiotropy or tended strongly towards this direction. It is noteworthy that another study focusing on local (rather than global) adaptation found the opposite relationship with increased pleiotropy for genes driving repeated local adaptation (49), using many of the same datasets and the same methods to identify repeated adaptation and pleiotropy.

**Fig. 4.**
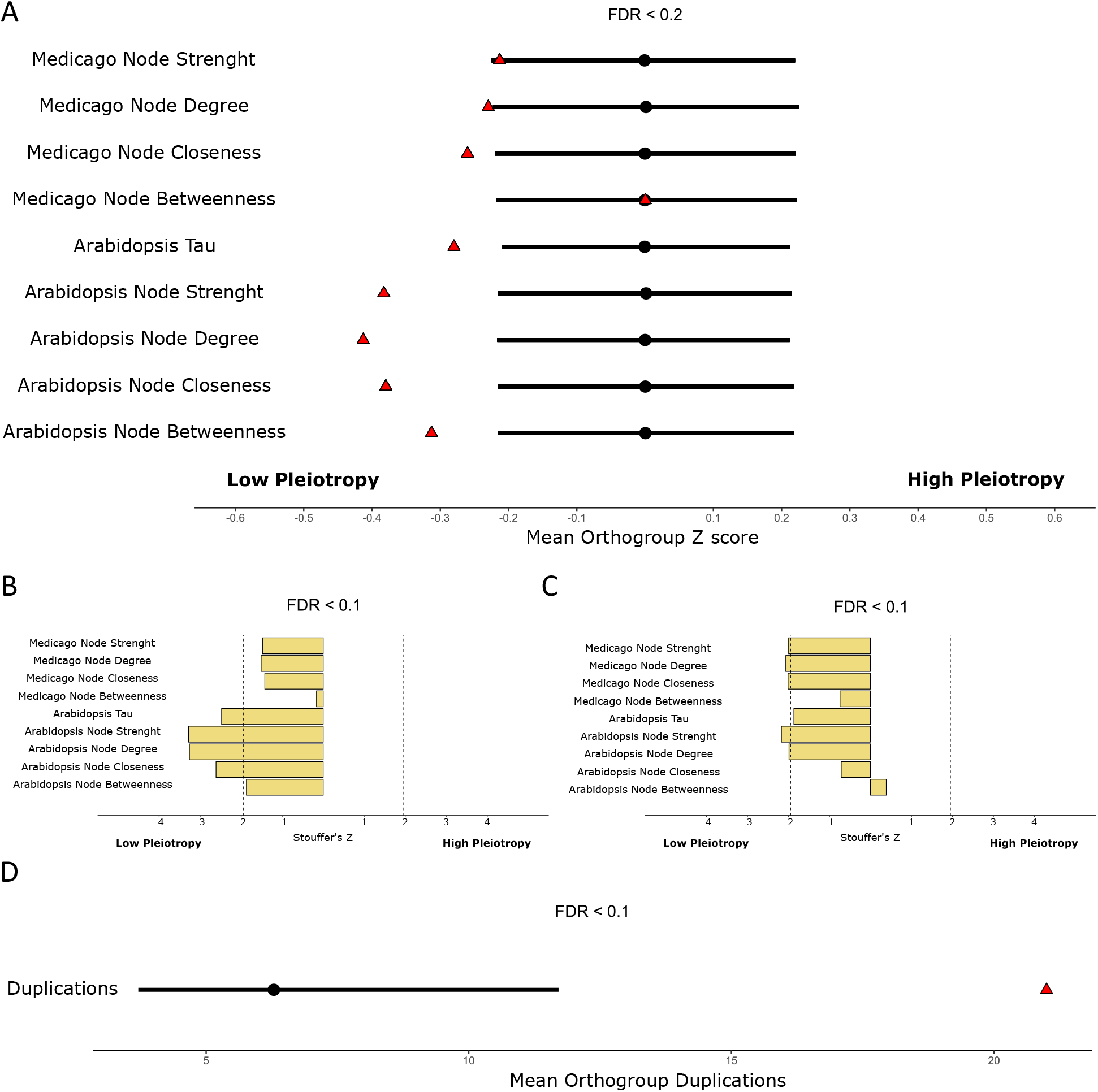
Orthogroups with repeated sweeps tend to be less pleiotropic and harbour more duplications. (A) Pleiotropy assessment of the relaxed set of 107 orthogroups with FDR < 0.2, against 10,000 random draws. Red triangles represent the average pleiotropy of the candidate set, while black circles represent the mean of 10,000 random draws. Black lines represent the 95% interval of the random draws means. Each row represents a different pleiotropy measure, labelled on the left. (B) Pleiotropy Stouffer’s Z of the top 33 orthogroups (FDR < 0.1), calculated using different pleiotropy measures labelled on the left. Dotted lines delimit the 95% confidence interval. (C) Pleiotropy Stouffer’s Z for the more conservative set of 15 orthogroups identified by intersecting the 33 candidate OGs of the main analysis (FDR < 0.1) with results derived from nine additional *PicMin* omitting closely related *Eucalyptus* and *Helianthus* species. Dotted lines delimit the 95% confidence interval. (D) Duplication bootstrapping assessment of the top 33 orthogroups (FDR < 0.1) against 10,000 random draws. The red triangle represents the average duplication number in the candidate set of orthogroups, while the black circle and line represent mean and 95% interval of 10,000 random draws respectively.

We further assessed pleiotropy within the more conservative set of 15 orthogroups identified by intersecting the 33 candidate OGs identified in the main analysis (FDR < 0.1) with results derived from nine additional *PicMin* omitting closely related *Eucalyptus* and *Helianthus* species. The correlation between global adaptation and decreased pleiotropy largely persisted (p < 0.05) when the assessment was restricted to this small set, indicating a robust association between genes driving global adaptation and decreased pleiotropy (Fig. 4C).

### Duplications are enriched in genes with repeated sweeps

Newly duplicated genes may experience selective sweeps due to positive selection if the resulting copies undergo sub- or neo-functionalization. Additionally, sub-functionalization reduces the pleiotropy of the original gene by distributing the function of the parent gene among new copies (85). This process may aid adaptation by easing the evolutionary constraints associated with a more pleiotropic ancestral gene (85,86). Considering this, we utilized the results obtained from *Orthofinder2* to investigate the involvement of gene duplications in recurrent global adaptation. We counted the number of paralogs within orthogroups as a representation of the number of duplication events and compared our candidates with repeated sweeps against a set of randomly chosen orthogroups, using the same statistical approach as the pleiotropy assessment. The 33 orthogroups with repeated sweeps (FDR < 0.1) exhibited a significant enrichment in gene duplications (p < 0.05), with a 3.3-fold-increase relative to the overall average for all orthogroups (Fig. 4D).

Next, we investigated whether the excess of duplicated genes observed in the recurrently globally adapted orthologous groups originated from small-scale single-copy gene duplications or from large-scale duplications due to polyploidization, as genes resulting from these two types of events evolve very differently (87,88). For this assessment, we retrieved lists of polyploidy-produced duplicated genes available for *Arabidopsis thaliana*, for which there is evidence of at least three ancient whole-genome duplication events (α, β and γ) (87,88,89).

Among the six driving genes identified in *A. thaliana* (*OmegaPlus* empirical p-values < 0.1 for the 33 *PicMin* orthogroups with FDR < 0.1, Fig. 3), two appear to have originated from whole-genome duplication events: ROXY1/ROXY2 (AT5G14070/OG0002218), which is dated back to the most recent duplication event α (86-14.5 mya), and a protein of unknown function (AT5G11000/OG0005857), which is dated to the β event (235-112 mya) (88,89). Extending this assessment to those orthogroups out of the 33 identified (*PicMin* FDR < 0.1) with a representative gene from *A. thaliana* (even though not necessarily an *A. thaliana* driving gene), we found that 9 genes in total (out of 22) appear to be polyploidy duplicated genes retained after one of the three ancient tetraploidy events (88,89). Although this enrichment does not achieve statistical significance (p > 0.05), it is clearly above the mean when assessed using a bootstrapping method akin to that employed for pleiotropy (SI Appendix, Fig. S9). Between the polyploidy duplicated genes with available functional annotation there were several other key developmental genes in addition to ROXY1/ROXY2, such as p300/CBP (OG0004755/ AT1G16705), ATIPT4 (OG0001035/AT4G24650), NPY (AT1G67900/OG0002632) and SLY1 (AT2G17980/OG0007322) as well as a few genes with key roles in the response to biotic stimuli, such as ERF19 (OG0006228/AT1G22810) and G-type lectin S-receptor-like serine/threonine-kinase (OG0002608/AT1G11340).

## Discussion

Here, we have uncovered evidence of 33 orthogroups (FDR < 0.1) with repeated global selective sweeps in multiple species (Fig. 3C). This discovery is noteworthy given the considerable evolutionary divergence among the analysed plant species and their diverse life history strategies and growth forms. These range from the herbaceous perennial monocot *P. hallii*, to eudicots encompassing both herbaceous annuals (*A. thaliana, C. rubella, M. truncatula, A. tuberculatus, Helianthus spp*.) and herbaceous perennials (*A. halleri*), to long-lived trees such as *Betula spp*., *Eucalyptus spp*., and *Populus spp*. (Fig. 1). While our study stands out for its focus on global adaptation and the exploration of more extensive phylogenetic distances, other studies have identified repeated signatures of local adaptation across independent populations and species residing in similar habitats or experiencing analogous selective pressures (32,33,34,35,36,37,38,39,40,41,49). Taken together, these studies suggest that while adaptation may commonly be polygenic and involve only subtle changes in allele frequency across many loci (90), it can also involve the contribution of key genes that appear to play particularly important roles, given their repeated involvement. In attempting to understand what determines the propensity of a gene to contribute to adaptation, it is important to consider both the biological characteristics of the gene and the influence of the methods used to detect selection.

Our set of genes with repeated sweeps includes many with demonstrated roles in abiotic and biotic stress responses in plants, as well as a few candidates with important functions related to growth and developmental processes (SI Appendix, Table S2). This echoes prior research based on rates of sequence substitution showing that protein function significantly influences these signatures of selection (91,92,93,94), as found in *Drosophila* (95,96), *Arabidopsis* (97), hominids (98), and other mammals (99,100). There is strong support for the hypothesis that host-pathogen interactions drive particularly rapid protein evolution, as immune and stress/defence response genes are often identified as targets of natural selection (91,92,93,94). Similarly, genes linked to defence responses, such as cytochrome P450 proteins (OG0001991), have been previously reported for their higher rates of sequence evolution (91,92). These findings are derived from methods based on high rates of nonsynonymous substitution that have very limited power to detect evolution that involves only a small number of impactful mutations and will therefore only tend to capture particularly strong and persistent selection pressure that drives many recurrent sweeps. By contrast, our sweep-based methods only detect more recent adaptation, as genomic signatures of selective sweeps degrade with time (14) but are able to detect the signatures arising from individual selective sweeps that may be missed by substitution-based methods. It is interesting that methods suited to detecting selection operating at very different timescales both tend to find genes involved in biotic interactions and defence as the targets of repeated natural selection.

### The role of gene duplication in adaptation

Gene duplication is thought to provide a source of genetic flexibility and facilitate of adaptation, through the means of sub and neo functionalization (85,86). Indeed, candidates for convergent local adaptation to temperature in two distantly related conifers, lodgepole pine (*Pinus contorta*) and interior spruce (*Picea glauca, Picea engelmannii, Picea glauca x Picea engelmannii*), were enriched for duplicated genes (34), but this was not found in the recent study of repeated local adaptation across plants (49). In this study we found a strong association between the orthogroups with signatures of repeated global adaptation and increased duplication number, with a 3.3x-fold increase relative to the average, despite our method penalizing duplicated genes with a conservative paralogue correction per orthogroup (Fig. 4D). However, we note that pleiotropy and number of duplications tend to be inversely correlated, therefore it can be tricky to separate the causative effects (SI Appendix, Table S3). Gene duplication may facilitate global adaptation by decreasing pleiotropy of a parent gene via sub-functionalization (85). Alternatively, if sub/neo-functionalization result in a new selective landscape for the novel copy of a gene, repeated sweeps would be expected as it evolves to improve this novel function. In either case, if the propensity for gene duplication is conserved over deep time, these mechanisms associated with duplication could partly explain our findings.

As polyploidy is a potential source of gene duplication, we also assessed whether the gene duplicates involved in repeated adaptation were derived from whole-genome or localized duplication. We found that a large portion of genes present in the repeated orthogroups (9 out of 22 genes in *A. thaliana*) may have originated during ancient polyploidization events, highlighting the important evolutionary role of whole-genome duplications in angiosperms. In *A. thaliana*, there is compelling evidence of at least three past tetraploidizations followed by subsequent re-diploidizations, with the oldest of these events dated before the moncot – dicot divergence, the second oldest dated before *A. thaliana* divergence from all the other dicots included in our study and the most recent event dated before the Brassicaceae radiation (89). During re-diploidizations some of the polyploidy-duplicated genes were retained and previous studies have shown that this process was highly biased towards certain gene functional classes, with some gene categories being expanded exclusively through these large-scale polyploidization–diploidization cycles (87,88). This is the case for several transcription factors and signal transducers involved in responses to biotic stimuli and secondary metabolisms, which are critical for survival (87,88), such as ERF19 (OG0006228/AT1G22810) and receptor-like protein kinases (RLKs) (OG0002608/AT1G11340), both identified here. Another category that has shown increased gene retention following whole-genome duplication events are developmental genes (87,88). Our analysis identified several *A. thaliana* genes belonging to this category present within the repeatedly globally selected orthogroups (*PicMin* FDR < 0.1), including ROXY1/ROXY2 (AT5G14070/OG0002218), p300/CBP (OG0004755/AT1G16705), ATIPT4 (OG0001035/AT4G24650), NPY (AT1G67900/OG0002632), and SLY1 (AT2G17980/OG0007322). Altogether, given that ancient whole genome duplications likely played a major role in the evolution of angiosperms by expanding certain gene functional classes to enhance growth and development as well as survival strategies against pathogens and other biotic threats (87,88,89), our results are consistent with an ongoing effect of this on the process of adaptation.

### Contrasting effects of pleiotropy in global vs. local adaptation

One characteristic of a gene that appears particularly important for adaptation is pleiotropy, where a single genetic locus influences multiple phenotypic traits (101). Pleiotropy is thought to constrain adaptation due to the possible detrimental effects of a mutational change affecting multiple biological processes (42,102). Fisher’s model of evolution provides a mathematical representation of this effect: the greater the number of trait dimensions, the lower the chance that a random mutation is beneficial, thus posing a reduced adaptive potential under higher pleiotropy (47). Similarly, if all genes have the same pleiotropy, larger mutations are more likely to incur negative fitness costs, due to the increased likelihood of overshooting the optimal trait value in some dimensions. Kimura (103) re-assessed Fisher’s model by incorporating the probability of fixation and concluded that mutations of intermediate effect would be most likely to contribute to adaptation. Later, Orr (104,105), considered the distribution of mutations fixed over an adaptive walk and found this would be approximately exponential, with alleles of larger effect fixing only earlier in the process, and later stages of evolution dominated by mutations of smaller effect, a prediction which has found empirical support in stickleback (106). This family of models therefore predicts that the rate of adaptation will be slower for organisms with more traits due to the greater amount of pleiotropy, and that most mutations driving adaptation will be of small effect (48).

Consistent with this “cost of complexity” hypothesis, genes and mutations with low pleiotropy have often been found as the target of parallel genetic and phenotypic evolution (42,102). Similarly, our research has pinpointed genes exhibiting low pleiotropy as recurrent targets of global adaptation across multiple plant species (Fig. 4). Furthermore, this evolutionary model has also found support in previous empirical studies, such as in yeast, where mutations affecting more phenotypic traits showed higher fitness costs hence implying a negative relationship between pleiotropic and fitness effects of mutations (70). Similarly, a study on Norwegian graylings populations showed that gene pleiotropy constrains both plastic and adaptive gene expression responses and highlighted the importance of genes with low pleiotropy in evolution (71). Taken together, these findings align with Fisher’s view of evolution, suggesting that functional changes that favour one process are more likely to have deleterious trade-offs on others if pleiotropy is high.

Further theoretical work on the importance of pleiotropy was motivated by empirical observations of substantial modularity (i.e. most genes affect few traits) along with larger per-trait mutation effects in more pleiotropic genes (107,108). These observations were incorporated into models showing that organisms with intermediate levels of pleiotropy would have the fastest adaptive rates, mitigating the cost of complexity (109). These species-level models have subsequently been interpreted as predicting the greatest contribution to adaptation within a species by genes with intermediate pleiotropy (73), which is reasonable but worthy of further directed theoretical study. Consistent with this prediction, empirical studies in *A. thaliana* (73), stickleback (32), ragweed (72), and *Heliconius* butterflies (74) have found evidence of intermediate pleiotropy driving repeated adaptation.

Another factor that can affect the impact of pleiotropy on adaptation is the spatial pattern of natural selection. If natural selection favours different trait optima in different locations of a species range (i.e., local adaptation), then migration will tend to counteract divergence, resulting in a general advantage for alleles of larger effect, as they can withstand the homogenizing effect of migration (6). To the extent that pleiotropic effects of a mutation are aligned with the direction of divergence in phenotypic optima, which seems common for modular traits (107,108), we would therefore expect more pleiotropic genes to contribute to local adaptation more readily, due to their larger effects overcoming the homogenizing effect of migration (109). By contrast, when a species adapts to a similar phenotypic optimum across its range (i.e., global adaptation), there is no tension with migration and therefore no additional advantage for alleles of larger effect (6). In accordance with this prediction (6,109), we found a strong association between decreased pleiotropy and genes involved in global adaptation (Fig. 4), which is also in line with Fisher’s and Orr’s models of evolution and with evidence of reduced pleiotropy in rapidly evolving adaptive genes reported in other studies (70,80,82,83). In further agreement with migration-selection theory predictions (6,109), the opposite pattern was found for genes driving local adaptation (i.e. increased pleiotropy) by another study using the same bioinformatic pipeline and statistical tests, and conducted on many of the same species, but studying signatures of local, rather than global adaptation (49). Taken together, these two studies provide the first controlled comparison of the genetics of repeated global vs. local adaptation. Observed differences in the pleiotropy of genes driving adaptive responses are consistent with population genetic predictions from models of spatially varying vs. spatially uniform selection. This therefore suggests that local adaptation may often look quite different than global adaptation, underlining the importance of considering the spatial pattern of selection when studying how evolution works.

## Materials & Methods

### Dataset selection

We downloaded raw sequencing data of 17 angiosperm plants and forest trees whole-genome sequencing (WGS) datasets from SRA and ENA (Fig. 1A and SI Appendix, Table S1).

The choice of WGS as the sole sequencing method for inclusion was imposed by the need for high quality dense SNP data by the software used in this study, *OmegaPlus* (45). Additionally, the datasets were chosen based on the following specific criteria: they encompassed natural populations in their native habitats, comprised non-invasive and non-domesticated species, included a minimum of 20 unrelated individuals sampled from five or more locations, and featured a high-quality reference genome, or one of a closely related species. We estimated each species natural range based on 1000 random observations downloaded from the Global Biodiversity Information Facility (http://www.gbif.org) using the *R* packages *rgbif* and *gbif*.*range* (110).

Finally, we explored the phylogenetic relationship between reference genome species using *TimeTree* (https://timetree.org/). Phylogenetic distances between species were calculated based on the *TimeTree* phylogeny using the R package *ape* (function: cophenetic).

### SNP calling

We applied a uniform SNP calling pipeline to all datasets for consistency. This pipeline was derived from a previous study and was selected as it optimizes the trade off between SNP quality and processing times (111). Raw *fastq* files were trimmed using *fastp* (112) with default settings. Clean reads were then aligned to reference genomes with *bwa-mem* (v0.7.17-r1188) (113), using 12 distinct reference genomes to map 17 datasets. If a species reference genome was not available, we used that of a closely related species. Three clusters of closely related species were mapped to the same reference genome (*B. pendula* and *B. platyphylla* mapped to *B. pendula*; *E. magnificata, E. sideroxylon* and *E. albens* to *E. grandis*; *H. annuus, H. petiolaris* and *H. argophyllus* to *H. annuus*). Following mapping, *samtools* was used to convert the alignment files from sequence alignment map (SAM) format to sorted, indexed binary alignment map (BAM) files, while discarding any alignment with mapping quality below 10 (-q 10) (114). The *MarkDuplicates* tool (115) from *Picard tools* was used to remove potential PCR duplicates and to set read groups. Indels were realigned using *GATK RealignerTargetCreator* followed by *GATK IndelRealigner* (116). After indel realignment, SNP calling was performed using *BCFTtools mpileup*, computing genotype likelihoods based on alignments with a minimum mapping quality of 5 (-q 5), followed by *BCFtools call* to identify single nucleotide polymorphisms (SNPs) from the pileup output and generate VCFs (114). Finally, we filtered raw VCF files with *VCFtools* (117) to retain only biallelic SNPs genotyped in at least 70% of the individuals, SNPs with quality value above 30 (--minQ 30), genotype quality above 20 (--minGQ 20) and minimum read depth above 5 (--minDP 5).

For downstream analyses, we retained SNP present at all minor allele frequencies except singletons. This retention of frequencies, often overlooked, was critical since the method employed to detect selective sweeps relies upon evaluating the patterns of linkage disequilibrium across genic regions (45). Therefore, filtering by allele frequency would introduce a bias by distorting LD patterns and could significantly decrease the detective power of the analysis, as excess of low frequency variants constitutes the main signature of a selective sweep (118). However, singletons are often bioinformatic artifacts and distinguishing them from real mutations can be challenging, hence their exclusion (119).

Finally, each dataset was filtered based on genomic relatedness (Fig. 1D). This step was taken to remove closely related individuals, as cryptic relatedness can potentially distort the estimation of regions under selection similarly to how it can confound GWAS (120). We used *Plink* (--genome function) to calculate relatedness (121), and the R package *PlinkQC* for filtering (122). *PlinkQC* aims to find the minimum number of individuals to remove to keep relatedness between any pairs below a chosen threshold (122). Individuals were systematically excluded from each species dataset to ensure that no relatedness scores between pairs exceeded 0.2, effectively removing any first and second-degree relatedness.

### Orthology inference

To assign genes to orthogroups, we first retrieved the amino acid sequences (proteomes) for each of the 12 reference species (Table S1). For each gene, we selected a primary transcript according to the longest isoform using custom scripts. Sequences were sorted by amino acid length, and each protein was given a name corresponding to the genomic coordinates of its gene. A Perl script was used to scan the proteins FASTA files. Upon encountering a duplicate sequence (i.e., a sequence with the same header), the script retained the first occurrence (the longest) and discarded subsequent occurrences. The decision is made based on whether the header has been encountered before. If it’s the first time encountering a particular header, the script writes both the header and sequence to the output file. Finally, *Orthofinder2* (51) was run with default settings using as input the 12 filtered proteomes, including only a single transcript per gene.

### Detection of selective sweeps using OmegaPlus

We used *OmegaPlus* (45) to scan each species dataset for global selective sweeps. *OmegaPlus* searches for specific LD patterns characteristic of recent hard selective sweeps and outputs the ω-statistic (13). LD-based methods have been shown to outperform SFS-based methods in the search for hard selective sweeps resulting in higher true positive rate (TPR), and *OmegaPlus* (45) has been consistently reported as the most sensitive tool to detect potential hard selective sweeps (10,15, 118). For our analysis of repeated sweeps across multiple species using *PicMin*, it is preferrable to use the most sensitive method at the cost of a higher false positive rate, in order to detect the highest number of true positive sweeps. This is not a reason of concern in comparative studies using *PicMin*, as the same false positives are unlikely to arise from independent analyses in different species (46). We extracted genes from each species’ VCF and added 1000 bp flanking regions on either side to include potential promoter and regulatory regions. Subsequently, *OmegaPlus* was run on each gene using a grid size of 3, resulting in 3 measurements: one at the first SNP, one at the last SNP and one equidistant between those 2 measurements. The minimum and maximum sizes of the subregion around a position that was included in the calculation of the ω-statistic were set to 500 and 100,000 base pairs, respectively. For each gene we retained the second scan, which was expected to lie approximately near the center of genes.

Following *OmegaPlus*, we ranked genes within each species by converting ω-scores to empirical p-values. An empirical p-value corresponds to the rank of a given gene’s selection score relative to all other genes from that species, therefore it reflects the strength of evidence against a null hypothesis of no selection (46). Gene empirical p-values were further summarized within each orthogroup by taking the lowest empirical p-value (i.e. the strongest selection evidence for a sweep) and applying a Dunn-Sidak correction to account for the number of paralogues within the orthogroup. This method of correcting for multiple comparisons within orthogroups only represents the contribution of the member gene with the strongest sweep signature within a given species (*i*.*e*. if two genes experience sweeps within an orthogroup in a given species, only the gene with the stronger signal contributes to the test).

We excluded orthogroups with more than 10 paralogues within a species, as they would be heavily penalized by the Dunn-Sidak correction and suffer from low power. Furthermore, we removed orthogroups with genes from less than 7 species from the analysis, as *PicMin* sensitivity would be reduced under such conditions. Finally, after these exclusions, we re-ranked the empirical p-values within each species to ensure uniform distributions of orthogroups empirical p-values for *PicMin*.

### Detection of selective sweeps using OmegaPlus: additional approaches

*OmegaPlus* was tested using the methodology outlined in the preceding section, with variations in the minimum window settings to explore robustness of our results. We experimented with both 200 and 1000 as minimum window sizes before ultimately opting for the intermediate value of 500. This was crucial, as research indicates that this setting can potentially significantly impact results and introduce increased stochasticity, particularly when employing very small minimum windows (118).

Additionally, we explored a genome-wide scan strategy, covering the entire genomes at intervals of 1000 bases. Subsequently, we extracted the scans falling within genes and aggregated these measurements on a per-gene basis by calculating the average for each gene, before applying the same methodology as before. However, we deemed this latter approach unsuitable due to its bias towards shorter genes, which tended to report more extreme average ω-scores.

Nevertheless, we evaluated the correlation between all different *OmegaPlus* runs per dataset and systematically examined the pleiotropy of the candidate orthogroups derived from each approach to enhance our confidence in the results.

### Testing for repeated global sweeps - PicMin

We used *PicMin* (46) to test for repetitive selective sweeps in 13,268 orthogroups across 17 species. *PicMin* uses order statistics to perform hypothesis tests on a set of ranked values, in this case empirical p-values derived from *OmegaPlus* ω scores, and identifies orthogroups enriched for large numbers of genes with low empirical p-values (46). Lower *OmegaPlus* empirical p-values correspond to higher ω scores, and indicate genes with stronger evidence of selection. For each orthogroup, *PicMin* provides a p-value representing the probability of generating ranks as extreme or more extreme than the observation under the null hypothesis of random genetic drift driving the *OmegaPlus* scores within each species. The method works as follows: for an orthogroup with genes in *n* species or lineages (17 in this case), under the null hypothesis the *n* empirical p-values representing the strength of evidence for selective sweeps in each species should follow a uniform distribution. If we order the empirical p-values within the orthogroup, the theory of order statistics shows they have marginal distributions that belong to the beta distribution, which *PicMin* uses to compute one-sided p-values. If the *x*^th^ rank in the list of ordered empirical p-values is low relative to the beta distribution, this indicates that the *x* species with the lowest empirical p-values all have stronger signatures of selective sweeps than would be expected by chance. Because some genes will always have low a rank within one species, we ignore the *x* = 1^st^ (lowest) ranked empirical p-value and consider all remaining higher ranks, effectively conducting tests of repeatability across two or more species. *PicMin* applies a multiple comparisons correction to the minimum p-value across all *x* = {2…*n*} contrasts, based on the methods by Tippett (123), Dunn and Sidak (124,125). This results in a final p-value that reflects the evidence that a particular orthogroup exhibits repeated adaptation. Finally, a multiple testing correction to account for the number of tested orthogroups was applied to the final per orthogroup p-values according to the Benjamini & Hochberg (1995) (126) formula, implemented in the R function *p*.*adjust* (method = “fdr”).

### Population structure assessment

For population structure assessment, SNPs with minor allele frequencies < 0.05 were discarded (*VCFtools*) (117) and each dataset was pruned by linkage disequilibrium (r^2^ >□0.4) using the *indep-pairphase* function of *Plink* in windows of 50 and step of 5 (121). Population structure was explored using principal component analysis (PCA) with *Plink* and ancestry inference with *fastSTRUCTURE* (127), testing Ks from 1 to 10. The representative admixture model for each dataset was determined using *fastSTRUCTURE* built-in *chooseK* function, which selects the model that maximises the log-marginal likelihood lower bound (LLBO) of the data and best explain strong population structure (127).

Weir & Cockerham (1984) *F*_ST_ (128) was calculated between the populations identified with *fastSTRUCTURE* on a per SNP site basis with the *vcftools* function *weir-fst-pop* (117). Within each species, individuals were assigned to populations according to the best K model Q coefficient, using as threshold of inclusion Q > 0.9. Average *F*_ST_ per gene was calculated by taking the mean across *F*_ST_ values of SNPs within each gene.

### Gene length

We assessed the association between gene length and *OmegaPlus* empirical p-values by calculating their Pearson correlation in each species, and examining the consistency of any patterning across species. We tested whether the mean gene length of the orthogroups with repeated sweep signatures identified with *PicMin* differed significantly from the expectation for randomly chosen genes. This was done by taking 10,000 random orthogroup draws of the same size as the significant set (33 orthogroups) and calculating the mean gene length of each draw. We then compared the mean of our candidate set against this null distribution.

### Recombination landscape

We tested the correlation between gene recombination rate and evidence for selective sweeps derived from the *OmegaPlus* analysis. To achieve this, we downloaded recombination rate data for *Arabidopsis thaliana* (129) and *Helianthus annuus* (130), corresponding to four distinct datasets in our analysis (*A. thaliana, H. annuus, H. petiolaris, H. argophyllus*). We then constructed density plots (in *R*) comparing *OmegaPlus* per-gene empirical p-values against the ranked average gene recombination rate.

Furthermore, we estimated recombination rates from site frequency spectrum (SFS) patterns for each dataset using the R package *FastEPRR* (131). The final filtered VCFs used for the main analysis were further refined to include only SNPs with minor allele frequency above 0.1 located on the major scaffolds (excluding short/fragmented scaffolds to avoid inaccurate phasing), and to retain no more than one SNP every 2500 bp, with *VCFtools* (117). This filtering step ensured a manageable SNP set for subsequent computations. Each species’ VCF was then split by chromosome and phased using *Beagle* (132). Recombination rates were estimated with *FastEPRR* (131) in non-overlapping 100,000 bp windows, and average per-gene recombination rates were calculated and then converted into empirical p-values in *R*. We used the R function *qbinom* to calculate the total number of driving genes expected to fall in either 5% tail of the distribution of genes’ recombination rates across all species.

### Nucleotide diversity

To determine if BGS could have influenced our repeatability results, we calculated genome-wide nucleotide diversity at each SNP site for each species using *VCFtools* (117). Next, we computed the average nucleotide diversity per gene for all genes in every species. We then compared the driving genes’ average nucleotide diversity relative to the average gene nucleotide diversity distribution within each species. Finally, we converted per-gene nucleotide diversity estimates into empirical p-values in *R*, and used the *R* function *qbinom* to calculate the total number of driving genes expected to fall in either 5% tail of the distribution of genes’ nucleotide diversity across all species.

### Estimation of pleiotropy

We estimated the amount of pleiotropy for each gene using two characteristics based on gene expression data: A) specificity of expression across tissues (77) and B) centrality within co-expression networks (78). To measure tissue specificity (A), we obtained *Arabidopsis thaliana* tissue expression data from Expression Atlas, accession *E-MTAB-7978* (75). This dataset includes tissue expression (transcripts per million - TPM) across developmental stages, tissue types and sub-tissue types. We computed the mean TPM across all developmental stages and sub-tissue types within each tissue type, to result in the mean TPM for each of the 23 tissue types. The tissue specificity metric τ was determined following the method by Yanai et al. (2005) (79) as:

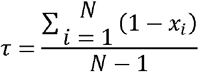

where, for a given gene, x_i_ corresponds to the mean TPM for a given tissue type normalised by the maximum mean TPM across N tissue types.

We converted τ scores into per-gene empirical p-values, treating higher τ estimates (corresponding to higher tissue specificity) as higher empirical p-values. We then applied the same methodology used to summarize *OmegaPlus* per-gene empirical p-values into orthogroups values, retaining the minimum empirical p-value (corresponding to minimum τ and therefore maximum pleiotropy) per orthogroup and applying a Dunn-Šidák correction to correct for the number of paralogues. Finally, we transformed per-orthogroup empirical p-values into Z-scores with a mean of 0 and standard deviation of 1 across all orthogroups.

This approach avoids assuming that specificity/pleiotropy is maintained across paralogues, which should be more representative than taking the mean τ per-orthogroup. In fact, calculating the mean τ per orthogroup significantly decreased the prevalence of high τ values in the genome-wide distribution. This suggests that tissue specificity within orthogroups varies significantly among paralogues, possibly due to neofunctionalization or sub functionalization. Additionally, we explored the same methodology in the reverse direction, considering lower τ estimates as lower empirical p-values. However, this alternative approach did not yield any substantial differences in the results.

To calculate the four centrality measures of co-expression networks (B), we built two networks using co-expression data from ATTED-II for *A. thaliana* and *M. truncatula*. The ATTED-II database summarises gene co-expression data derived from RNA-seq and microarray sources in a condition-independent manner, which is given as a standardised z-score between a given pair of genes (76). To construct co-expression networks, genes were treated as nodes and z-scores as edges, where positive z-scores denote positive co-expression and vice-versa for negative z-scores. Co-expression gene tables were downloaded for both species: *A. thaliana* = Ath-u.c3-0 and *M. truncatula* = Mtr-u.c3-0. We discarded all edges with -5 < z < 2.33 following the recommendations for significant negative/positive co-expression.

The *A. thaliana network* included 18,570 genes, while *M. truncatula* network included 17,786 genes. Networks were generated using the *igraph* package in R. Node betweenness and closeness were calculated using the *estimate_betweenness()* and *closeness()* functions respectively. Node degree and strength were calculated as the number of edges and the sum of all edge absolute z-scores respectively. We then condensed the resulting gene centrality measures into per-orthogroup Z-scores with mean 0 and standard deviation of 1 across all orthogroups, with the same approach used for tissue specificity τ scores and testing both directions for the initial conversion of centrality scores into empirical p-values.

To test whether genes with repeated sweep signatures had high/low values of pleiotropy, for both the tissue specificity (A) and centrality measures (B) we used a bootstrapping method to compare their values to those of randomly chosen genes. For each measure, we calculated the mean z-score of the candidate set for repeatability, including 33 OGs. We then performed 10,000 random draws, each comprising the same number of orthogroups as the candidate set and calculated the mean of each random draw. Finally, we assessed whether the mean of the candidate set fell within the 95% confidence interval of the means from the10,000 random draws. For the smaller sets of candidates, we assessed pleiotropy by calculating Stouffer’s Z score (133) for tissue specificity and centrality measures using the following formula:

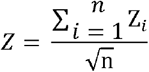

### Gene duplication

We used the output from *Orthofinder2* to count duplication events within orthogroups, including both terminal and nonterminal nodes. We then used this data to test whether the candidate set with repeated sweep signatures was significantly enriched for duplications, using the same bootstrapping method used for pleiotropy (described in the previous section).

## Data Availability

The scripts for SNP calling are available at: https://github.com/GabrieleNocchi/snp_calling_bcftools_slurm

The scripts for the population structure analysis are available at: https://github.com/GabrieleNocchi/population_structure_analysis

The scripts for the main analyses are available at: https://github.com/GabrieleNocchi/RepSweeps

References and links to the genomic resources for each dataset are available in SI Appendix, Table S1.

## Acknowledgements

Funding was provided by NSERC Discovery and Alberta Innovates, with computational resources and support provided by the Digital Research Alliance of Canada.

## Notes

### Competing Interest Statement

The authors have declared no competing interest.

### Summary of Updates

Added an extra test in sup mat in regards to duplication occurred after WGD. Corrected a minor error in the description of the recombination rate calculation method.

