## Supplementary material for "Repeated global adaptation across plant species": SI Appendix

PNAS

### Supporting Information for
Repeated global adaptation across plant species

Gabriele Nocchi[1], James R. Whiting[1], Samuel Yeaman[1]

1. Department of Biological Sciences, University of Calgary, Calgary, Canada T2N 1N4

Corresponding author: Gabriele Nocchi

**This PDF file includes:**

Supporting Text
Figures S1 to S21
Tables S1 to S4
SI References

### Supporting Text

##### Possible biases and artifacts

*PicMin* identifies repeated orthogroups based on statistical support across different species or lineages (1). To test *PicMin's* robustness to unexpected biases, we performed 1000 additional *PicMin* analyses, each time randomly shuffling the *OmegaPlus* empirical p-values within each species. After correcting for multiple tests, the mean number of orthogroups with FDR < 0.1 per-permutation was 0.024. These 24 orthogroups with FDR < 0.1 arose within 9 out of the 1000 permutations (SI Appendix, Fig. S10). Generally, the *PicMin* analyses based on permuted data reported few orthogroups with false discovery rates below 0.9 compared to the actual *PicMin* analysis, and rarely any below 0.1 (SI Appendix, Fig. S10 and Fig. S11). The FDR distributions derived from *PicMin* based on the shuffled data appear clearly different from that of the actual data, with the latter exhibiting a markedly longer left tail (SI Appendix, Fig. S12).

As *PicMin* identifies orthogroups with consistently extreme signatures across multiple species, any source of bias arising from some characteristic of a gene can also bias *PicMin* if the characteristic tends to be conserved over deep time. Gene length is one such characteristic that might contribute to bias, based on how we ran *OmegaPlus* using one scan near the centre of each gene. This analysis choice might lead to reduced detection power in longer genes, as the signal of a selective sweep near a gene edge would be more attenuated at the middle of the gene due to the more extensive decay of linkage disequilibrium, compared to shorter genes. Consequently, *OmegaPlus* might exhibit reduced sensitivity for larger genes. Our investigation revealed that this concern was unfounded, as there were generally low correlations between empirical p-values for the sweep signatures and gene length within each species (SI Appendix, Fig. S13). No consistent pattern was observed in the relationship between gene length and empirical p-value across species, as 10 species had $r > 0$, while 7 had $r < 0$; (SI Appendix, Fig. S14). Furthermore, when assessing the average gene length in the 33 orthogroups candidates for repeated global adaptation (FDR < 0.1), we found no significant difference in mean gene length compared to random expectations. If anything, our candidate orthogroups exhibit a tendency toward increased gene length (SI Appendix, Fig. S15), counter to the expectation favouring shorter genes under this type of bias.

To strengthen our confidence in the results, we next assessed the candidates derived from each of three additional approaches to implementing sweep scans using *OmegaPlus*, with variations in settings (refer to *Methods* for details). All three approaches showed high correlations between the resulting *OmegaPlus* empirical p-values and the empirical p-values obtained with the main approach implemented above (*OmegaPlus* with minimum window of 500 bp) (SI Appendix, Table S4). Remarkably, all approaches yielded consistent results in the association with pleiotropy, highlighting a robust relationship between global adaptation and low pleiotropy (SI Appendix, Fig. S16). While our analysis of pleiotropy assumes that estimates of tissue specificity from *A. thaliana* are representative of patterns in other species, specificity in expression has been shown to decline slowly among orthologues (2). In addition, our findings are robust to different measures of pleiotropy (Fig. 4).

Another potential source of bias is in the recombination landscape around a gene, as regions with lower recombination rates may harbour higher LD and lower diversity (3), which could lead to an enrichment of low *OmegaPlus* empirical p-values in these regions. It is also possible that sweep signatures may take longer to degrade in regions of low recombination. However, recombination events play a crucial role in generating the typical selective sweep linkage disequilibrium patterns recognized by *OmegaPlus*, therefore recombination must be substantial to obtain higher ω scores (4,5). This was largely reflected in our assessment based on available recombination data (see *Methods*), which showed higher density of lower *OmegaPlus* empirical p-values in regions with moderate to high recombination, with the exception of *H. argophyllus* (SI Appendix, Fig. S17).

Overall, *OmegaPlus* demonstrated robustness to misidentifying regions with lower recombination rates as sweeps (SI Appendix, Fig. S17). In addition, we estimated average gene recombination rates de novo for each species based on the site frequency spectrum (SFS) (see *Methods* for details). The average recombination rates of the driving genes (*OmegaPlus* emp-p < 0.1) of the 33 orthogroups with repeated signals of global selection identified with *PicMin* (FDR < 0.1) showed significant variability and no consistent pattern (SI Appendix, Fig. S18). If the recombination rates of the driving genes are randomly distributed relative to the background genomic rates, we would expect up to 13 driving genes in total to fall within either 5% tail of the distribution of genes recombination rates within species (95% confidence limit of a binomial distribution, based on 160 driving genes with available estimated recombination rate across all species). Our results do not significantly exceed this expectation, as we found 11 driving genes in the top 5% of the recombination rate distribution within species and 12 in the bottom 5% (SI Appendix, Fig. S19).

Finally, even though a previous study has shown that background selection (BGS) does not generate the LD signature typical of selective sweeps and does not tend to cause any deviation in *OmegaPlus* ω scores relative to neutrality (6), we assessed whether BGS could have driven our repeatability results. Using nucleotide diversity as a proxy for assessing BGS, we found that the driving genes of the 33 orthogroups identified with *PicMin* exhibit significant variability in nucleotide diversity, showing no consistent pattern, indicating the absence of this potential bias (SI Appendix, Fig. S20). If the nucleotide diversity of the driving genes is randomly distributed relative to the background gene diversity, we would expect up to 14 driving genes in total to fall within either 5% tail of the distribution of genes nucleotide diversity within species (95% confidence limit of a binomial distribution, based on 172 driving genes across all species). Our results do not significantly exceed this expectation, as we found eight driving genes in the top 5% of the recombination rate distribution within species and four in the bottom 5% (SI Appendix, Fig. S21).

### Supporting Figures

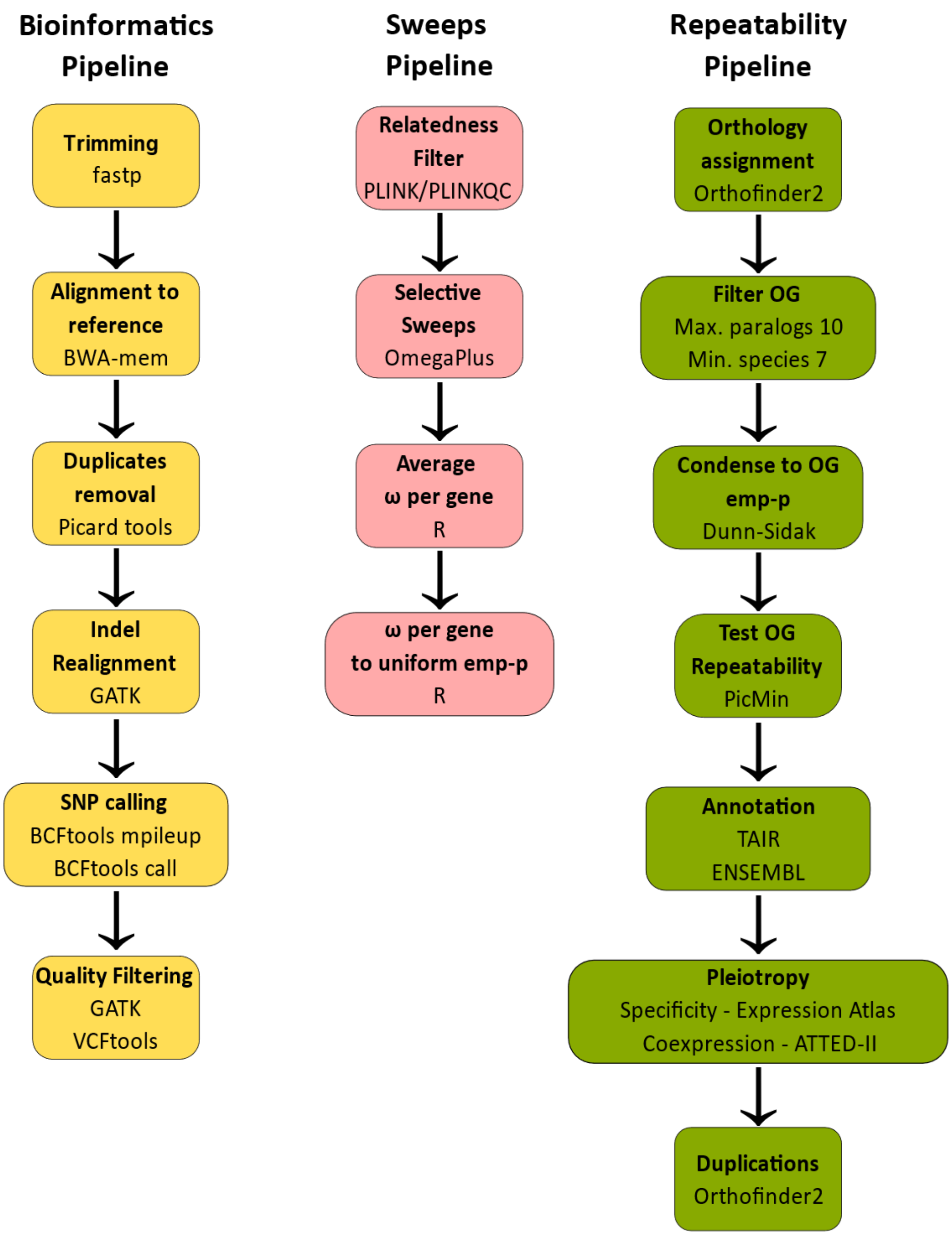


**Fig. S1.** Schematic summary of methods. The bioinformatics pipeline used for SNP calling in yellow, the selective sweep detection pipeline in pink and the repeatability and pleiotropy/duplications assessment pipeline in green.

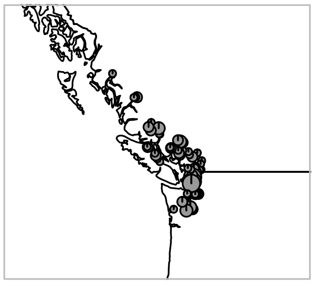

*P. trichocarpa*

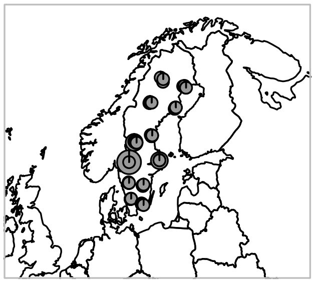

*P. tremula*

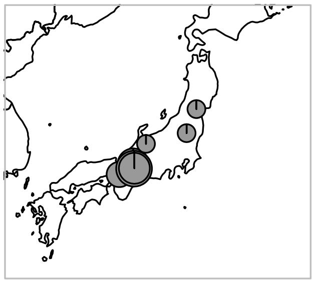

*A. halleri*

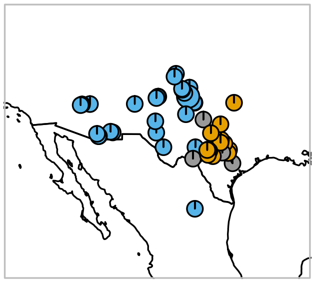

*P. hallii*

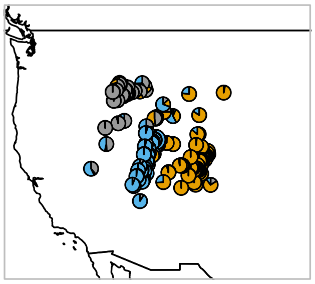

*P. stricta*

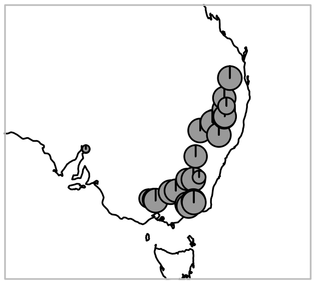

*E. albens*

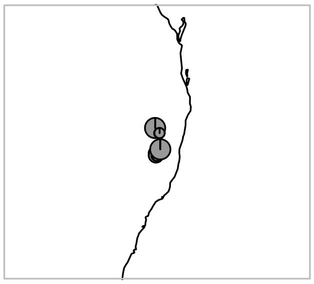

*E. magnificata*

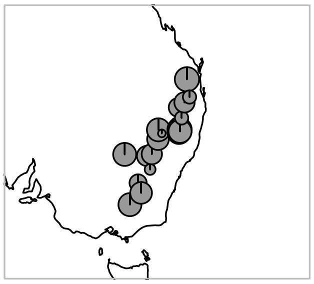

*E. sideroxylon*

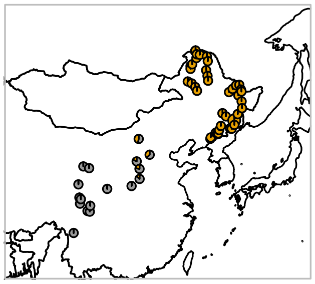

*B. platyphylla*

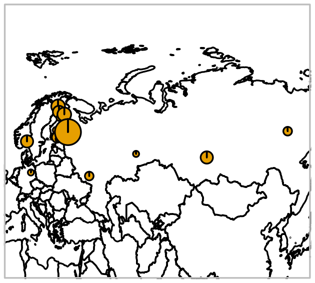

*B. pendula*

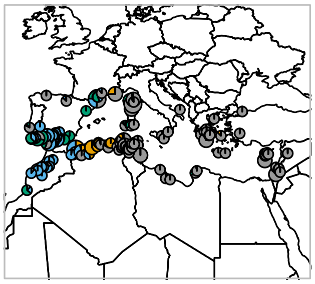

*M. truncatula*

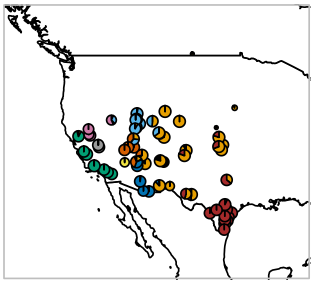

*H. annuus*

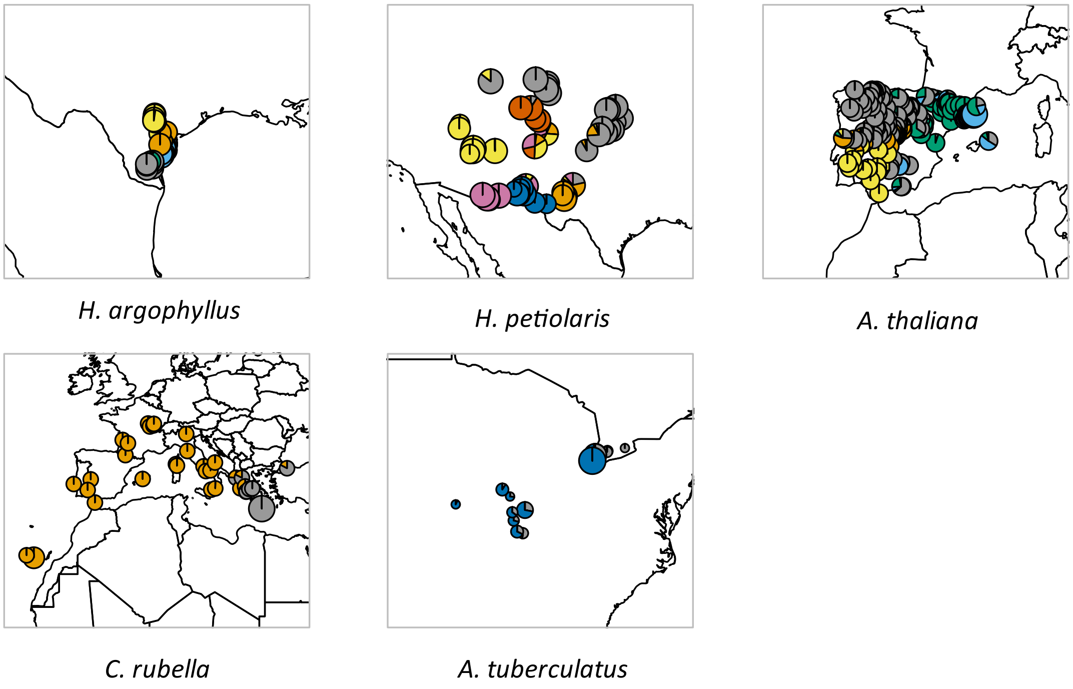


**Fig. S2.** *fastSTRUCTURE*[7] ancestry pie plots of the 17 datasets included in this study. The size of the pies is proportional to the number of individuals at each sampling location. The number of colors in each plot reflects the K value of the model that maximises the log-marginal likelihood lower bound (LLBO) of each dataset and best explains strong population structure, according to *fastSTRUCTURE*[7].

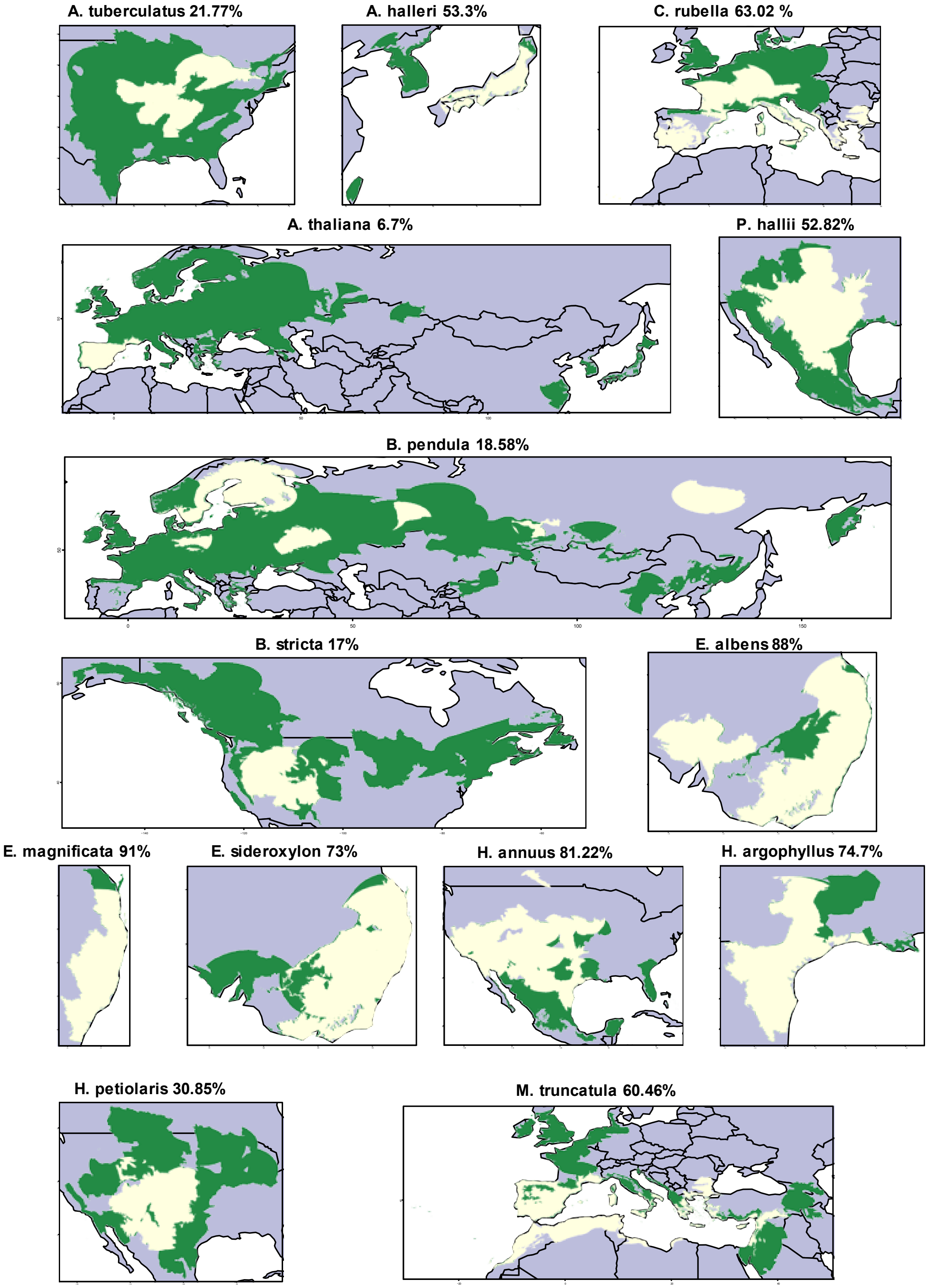
A. tuberculatus 21.77%
A. halleri 53.3%
C. rubella 63.02 %
A. thaliana 6.7%
P. hallii 52.82%
B. pendula 18.58%
B. stricta 17%
E. albens 88%
E. magnificata 91%
E. sideroxylon 73%
H. annuus 81.22%
H. argophyllus 74.7%
H. petiolaris 30.85%
M. truncatula 60.46%

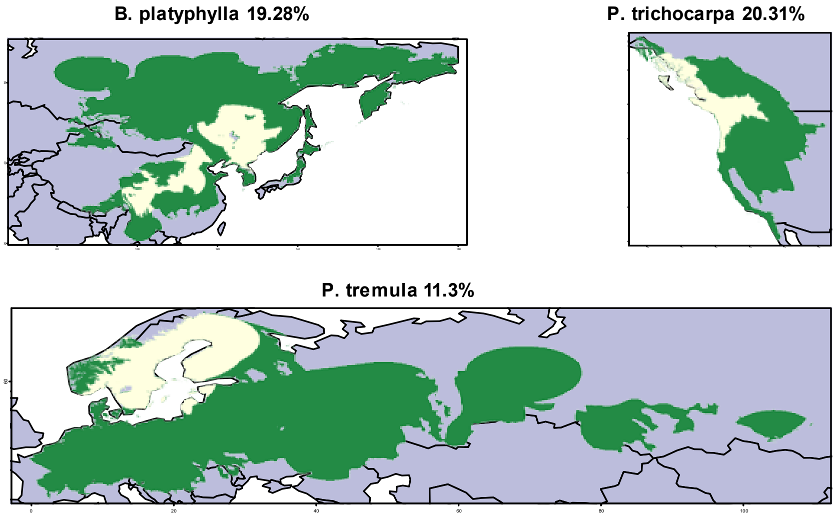


**Fig. S3.** Datasets coverage of species ranges. The maps illustrate the sampling coverage of each dataset (light yellow) over the estimated native range of species (light yellow + green), based on *GBIF* observations (http://www.gbif.org). The approximate coverage (%) is reported above each map.

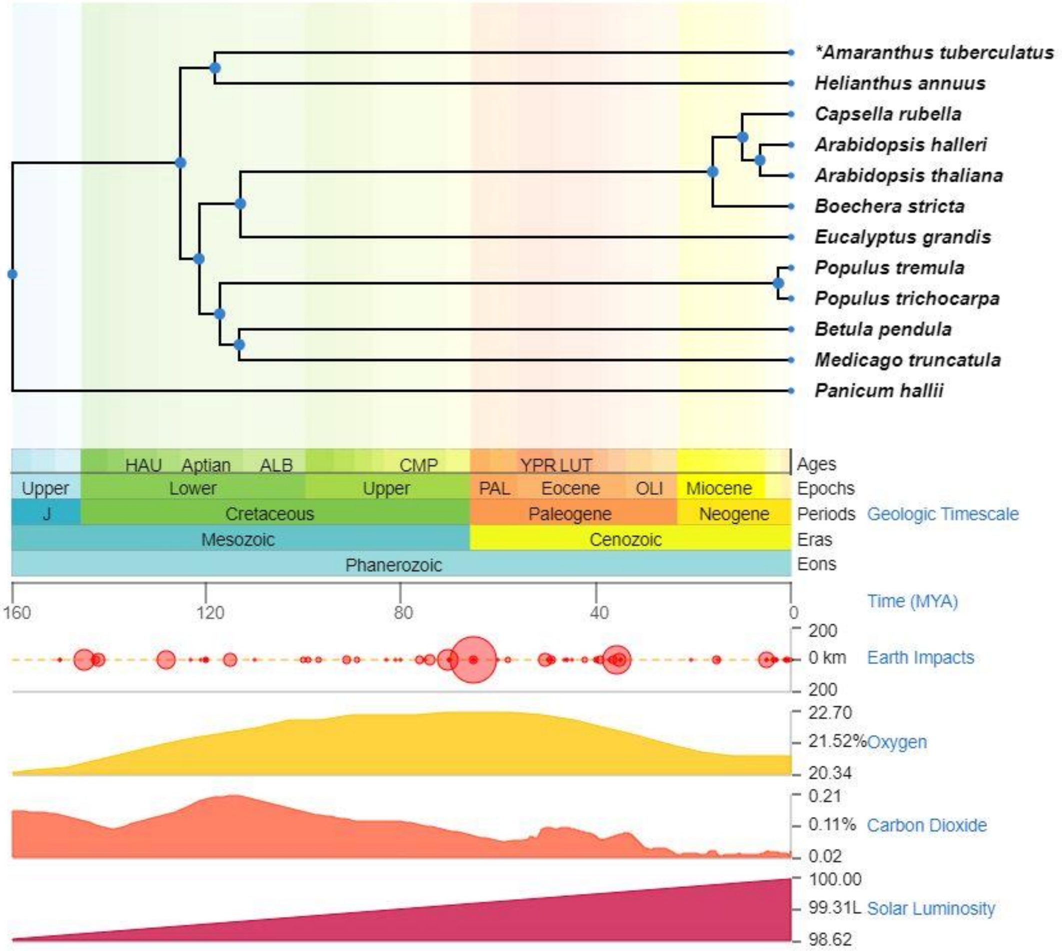


**Fig. S4.** *TimeTree*[8] time-calibrated phylogeny of the 12 reference genome species analyzed in this study. Asterisks indicate instances where *TimeTree* opted for a substitute species due to the unavailability of data on the target species. *Amaranthus tuberculatus* was replaced with *Amaranthus hybridus*.

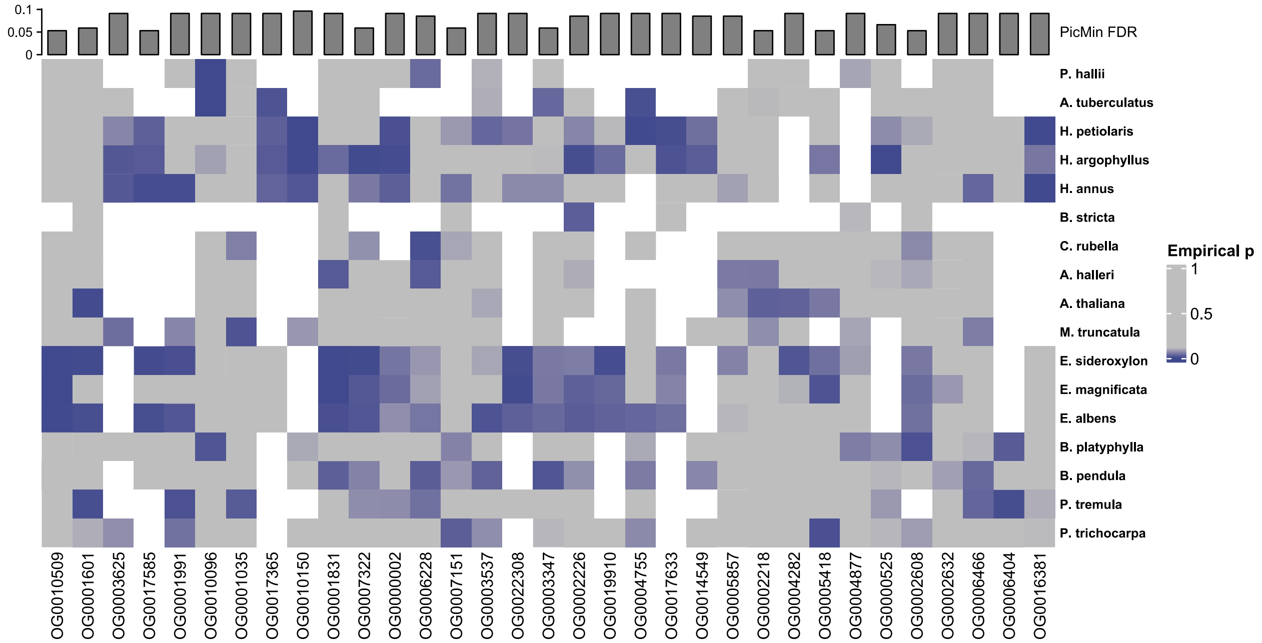


**Fig. S5.** Heatmap of *OmegaPlus*[9] empirical p-values of the 33 orthogroups with *PicMin*[1] FDR < 0.1. Genes with *OmegaPlus*[9] empirical p-value < 0.1 (driving genes) are highlighted using a blue gradient. *OmegaPlus*[9] empirical p-values > 0.1 are colored in grey. Species are ordered by phylogenetic distance along the Y axis. A white cell indicates that an orthogroup was not tested in a species.

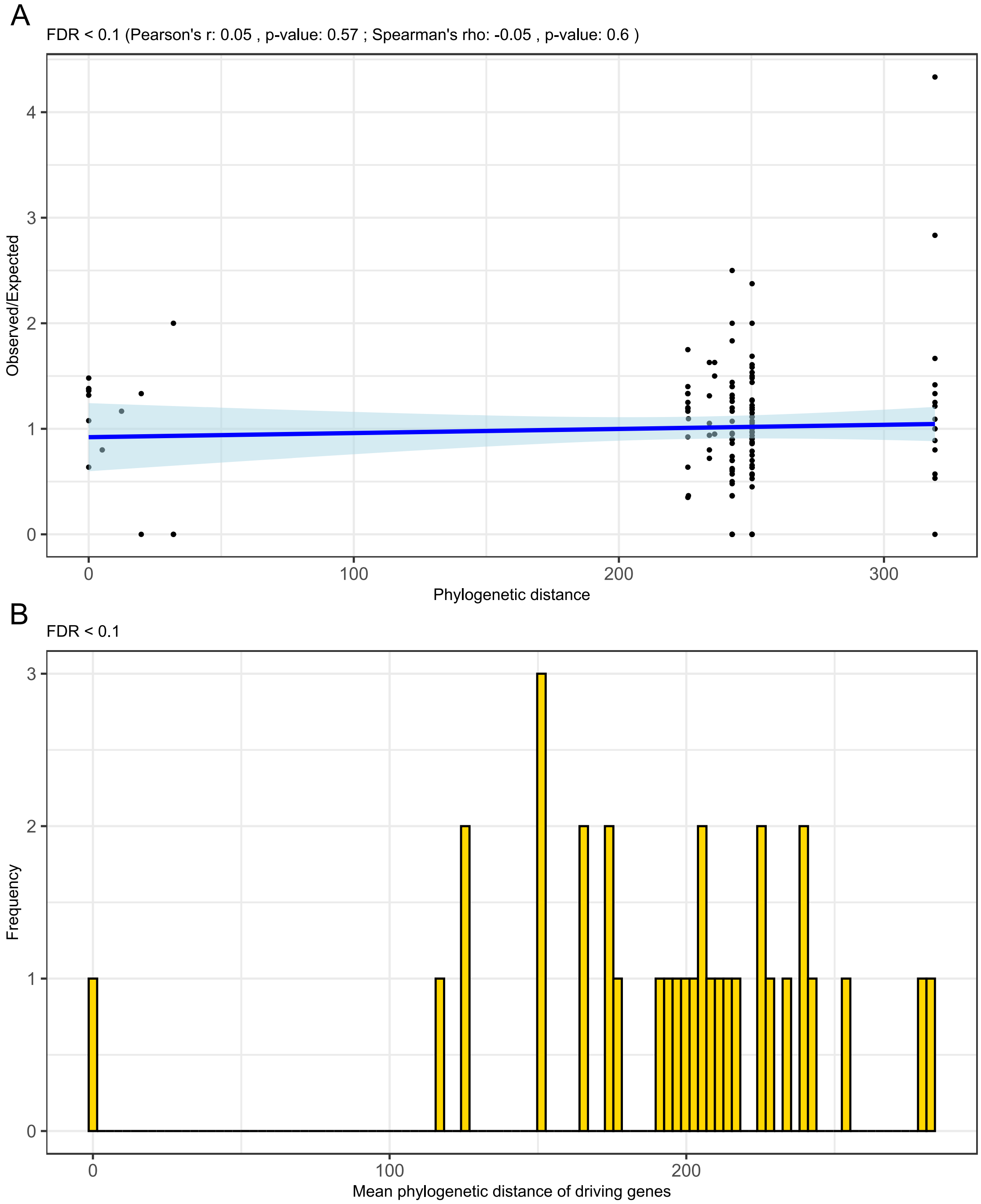


**Fig. S6.** Phylogenetic distance assessment. (A) Correlation between phylogenetic distance between each possible pair of species and the ratio of observed versus expected driving genes overlap between pairs of species, according to the expectation of a hypergeometric distribution. The blue line and light blue shading represent the regression line and its 0.95 confidence interval respectively. (B) Distribution of the mean phylogenetic distance between the driving genes (*OmegaPlus*[9] empirical p-value < 0.1) of the top 33 orthogroups (*PicMin*[1] FDR < 0.1).

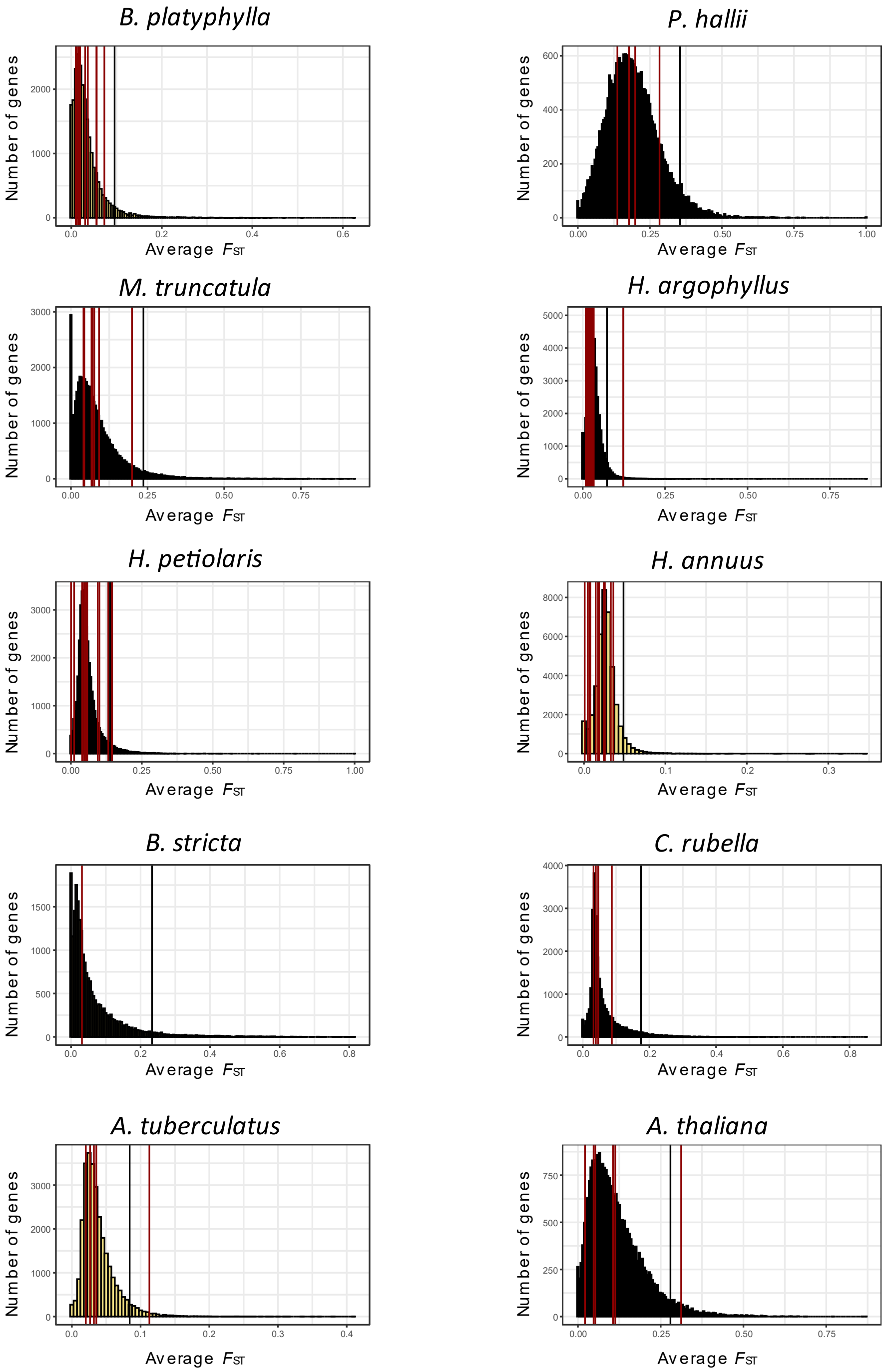


**Fig. S7.** Genes $F_{ST}$[10] assessment in each species. Red lines mark driving genes $F_{ST}$. Yellow histograms represent the genes $F_{ST}$ distribution within each species. Black bars represent the 95% interval upper-bound of the genes $F_{ST}$ distribution.

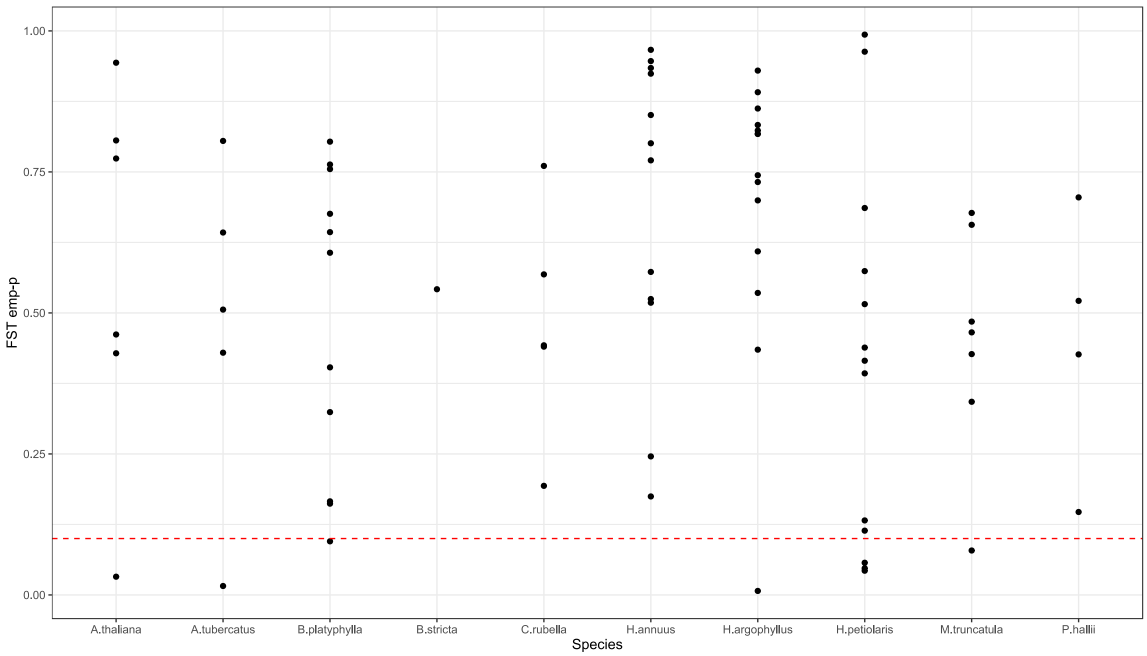


**Fig. S8.** $F_{ST}$[10] assessment in each species. The Y axis represents the $F_{ST}$ empirical p-value or rank, with smaller empirical p-values corresponding to higher $F_{ST}$. Points represents driving genes in each species (*OmegaPlus*[9] empirical p-values < 0.1 in the *PicMin*[1] FDR < 0.1 orthogroups). The red dashed line corresponds to an empirical p-value of 0.1.

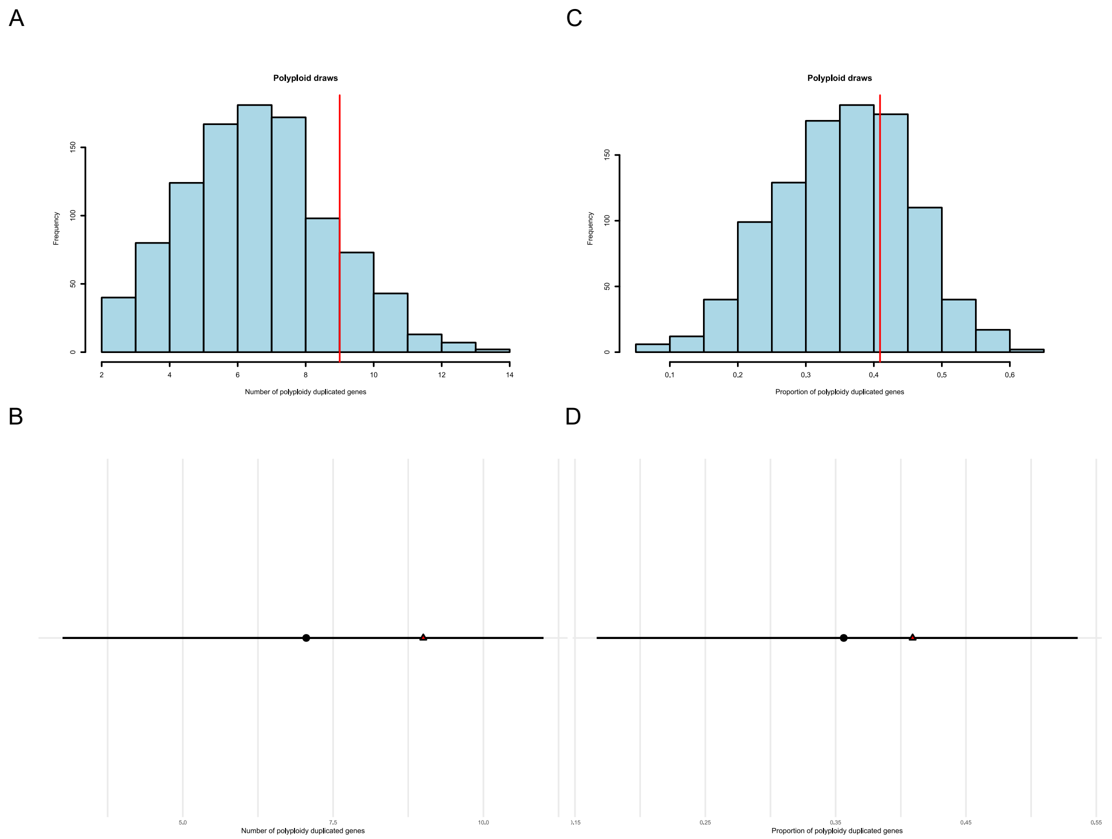


**Fig. S9.** Polyploidy duplicated genes assessment. (A) Histogram of the number of *A. thaliana* polyploidy duplicated genes in 1000 random draws, each draw composed of 22 *A. thaliana* orthogroups. The red line represents the number of polyploidy duplicated genes (9) in the 22 candidate *A. thaliana* orthogroups (*PicMin*[1] FDR < 0.1). (B) Polyploidy duplicated genes bootstrapping assessment. The red triangle represents the number of polyploidy duplicated genes (9) in the 22 candidate *A. thaliana* orthogroups (*PicMin*[1] FDR < 0.1). The black circle represents the mean number of *A. thaliana* polyploidy duplicated genes per draw in 1000 random draws (each draw composed of 22 random *A. thaliana* orthogroups). (C) Histogram of the proportion of *A. thaliana* polyploidy duplicated genes in 1000 random draws, each draw composed of 33 random orthogroups (including all species). The proportion of polyploidy duplicated genes was calculated out of the number of *A. thaliana* orthogroups present in each draw. The red line represents the proportion of polyploidy duplicated genes (9 out of 22) in the 33 candidate orthogroups (*PicMin*[1] FDR < 0.1). (D) Polyploidy duplicated genes bootstrapping assessment. The red triangle represents the proportion of *A. thaliana* polyploidy duplicated genes (9 out of 22) in the 33 candidate orthogroups (*PicMin*[1] FDR < 0.1). The black circle represents the mean proportion of *A. thaliana* polyploidy duplicated genes per draw in 1000 random draws (each composed of 33 random orthogroups). The proportion of polyploidy duplicated genes was calculated out of the number of *A. thaliana* orthogroups present in each draw. The black lines in plots (B) and (D) represent the 95% confidence intervals for the mean of the random draws.

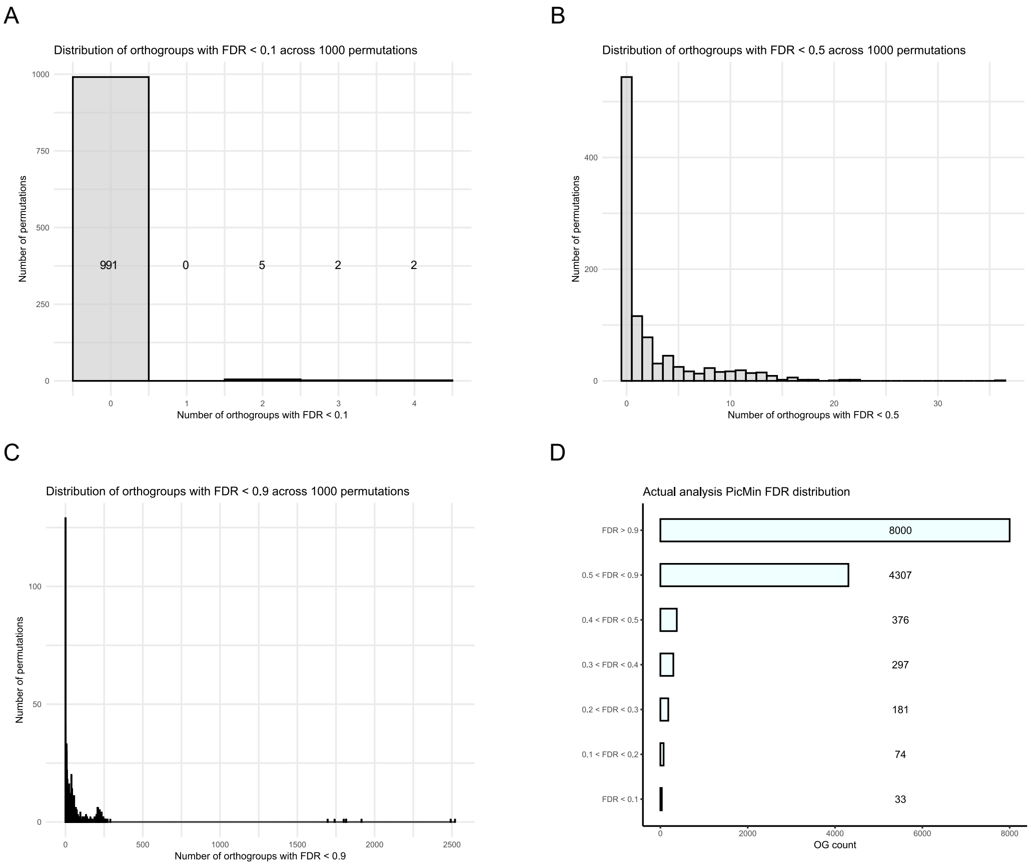


**Fig. S10.** 1000 *PicMin*[1] permutations results. (A) Distribution of *PicMin*[1] permutations tests reporting orthogroups with FDR < 0.1 (B) Distribution of *PicMin*[1] permutations tests reporting orthogroups with FDR < 0.5 (C) Distribution of *PicMin*[1] permutations tests reporting orthogroups with FDR < 0.9 (D) Count of orthogroups across FDR windows in the actual *PicMin*[1] analysis.

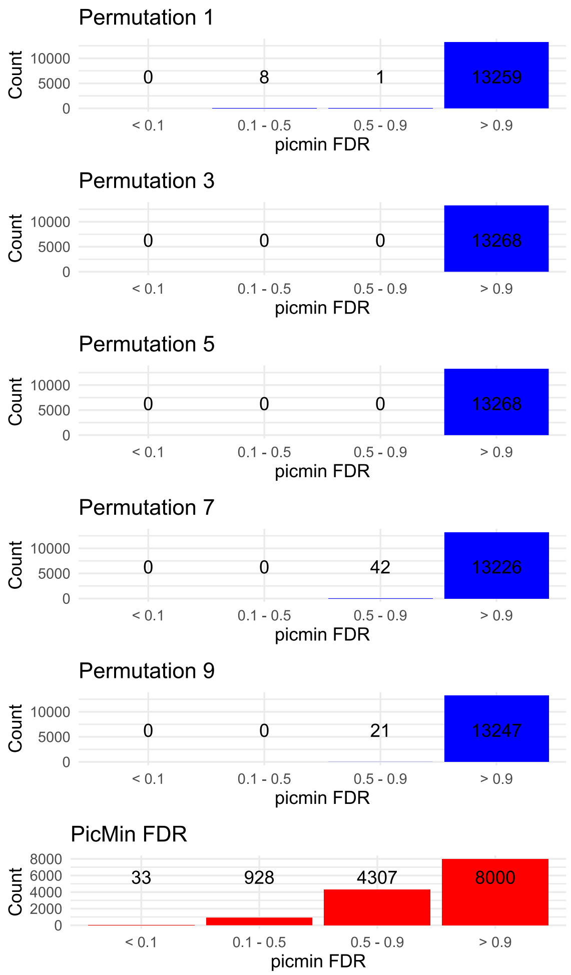


**Fig. S11.** *PicMin*[1] permutations FDR count. Bar plots displaying the count of orthogroups across four FDR windows ( < 0.1, 0.1 – 0.5, 0.5 – 0.9, > 0.9) for 10 *PicMin*[1] tests (out of 1000) based on data permutations (in blue) against *PicMin*[1] on the actual data (in red).

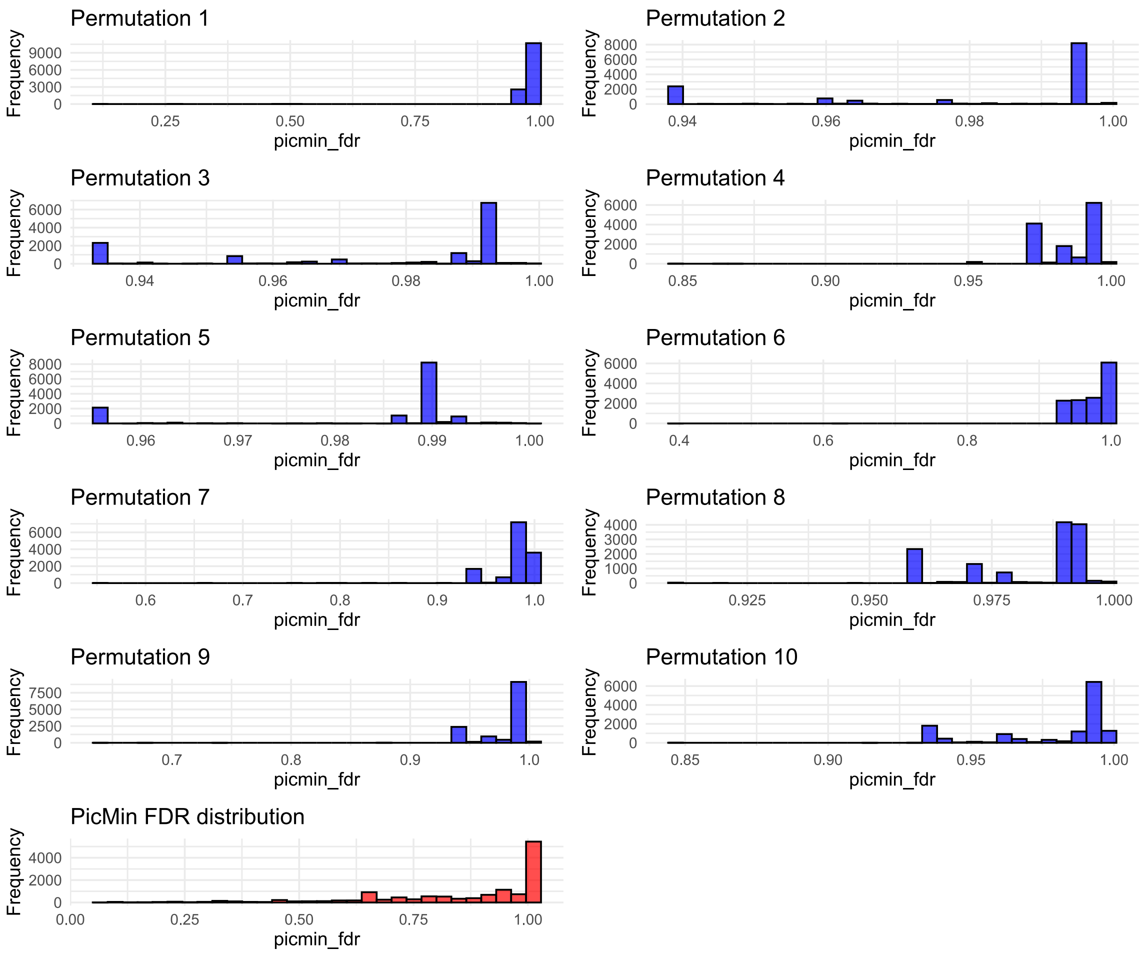


**Fig. S12.** *PicMin*¹ permutations FDR distribution. Distribution of *PicMin*¹ FDR for 10 *PicMin*¹ tests (out of 1000) based on data permutations (in blue) against the *PicMin*¹ FDR distribution of the actual data (in red).

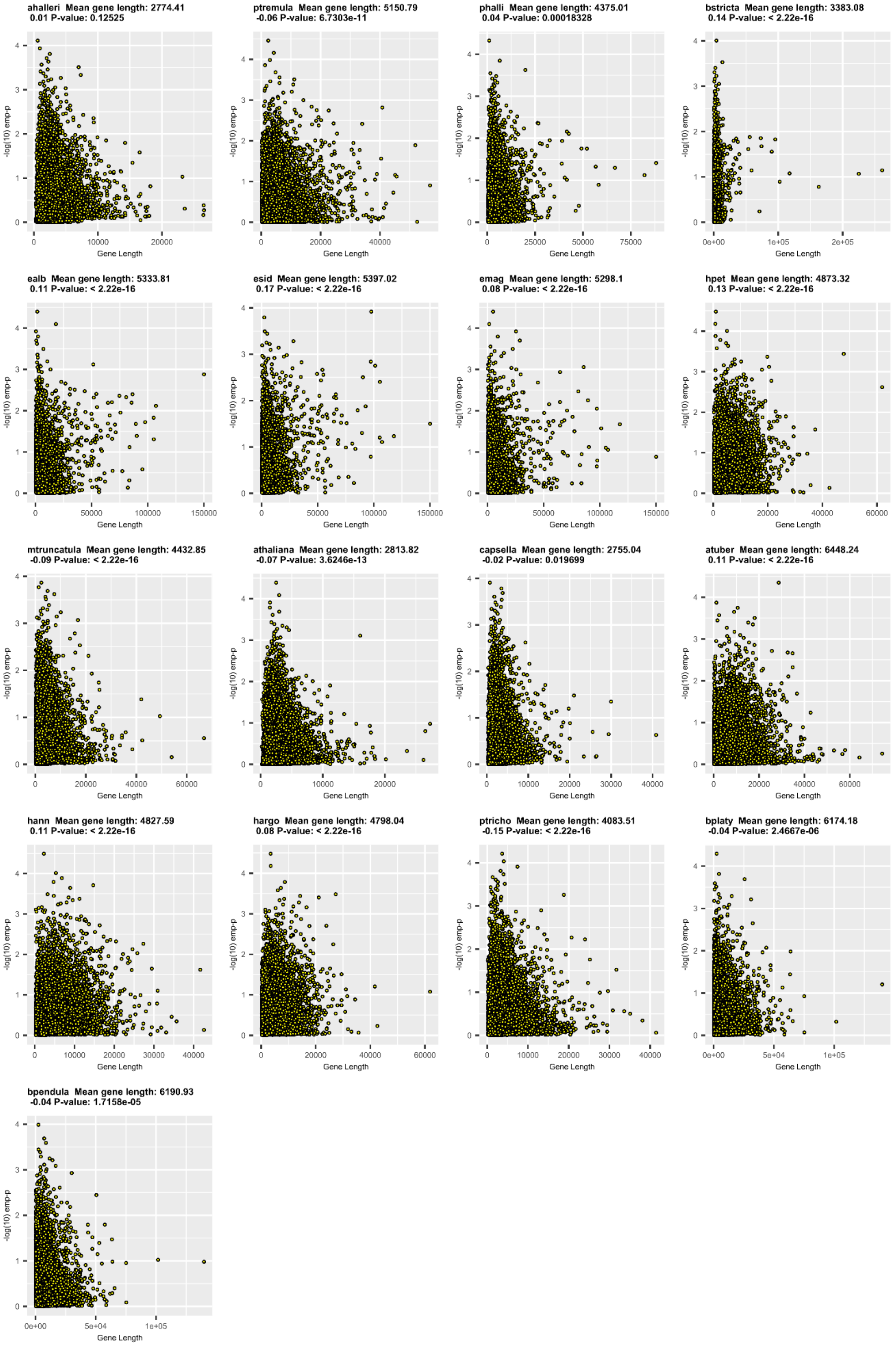


**Fig. S13.** Correlation between gene length and *OmegaPlus*[9] empirical p-values (-log10) in each species. Species abbreviated name, mean gene length, *Pearson's r* and correlation test p-value are given above each plot.

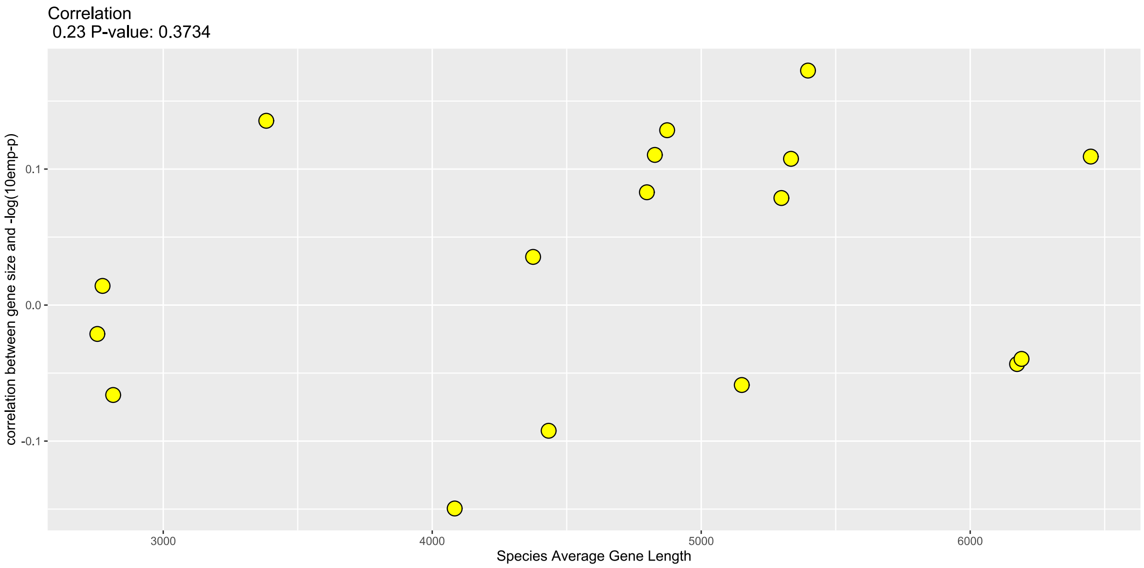


**Fig. S14.** Correlation between species average gene length and the Pearson's r of the correlation between gene size and *OmegaPlus*[9] empirical p-values (-log10) (Fig. S8). Overall *Pearson's r* and correlation test p-value given on top of plot.

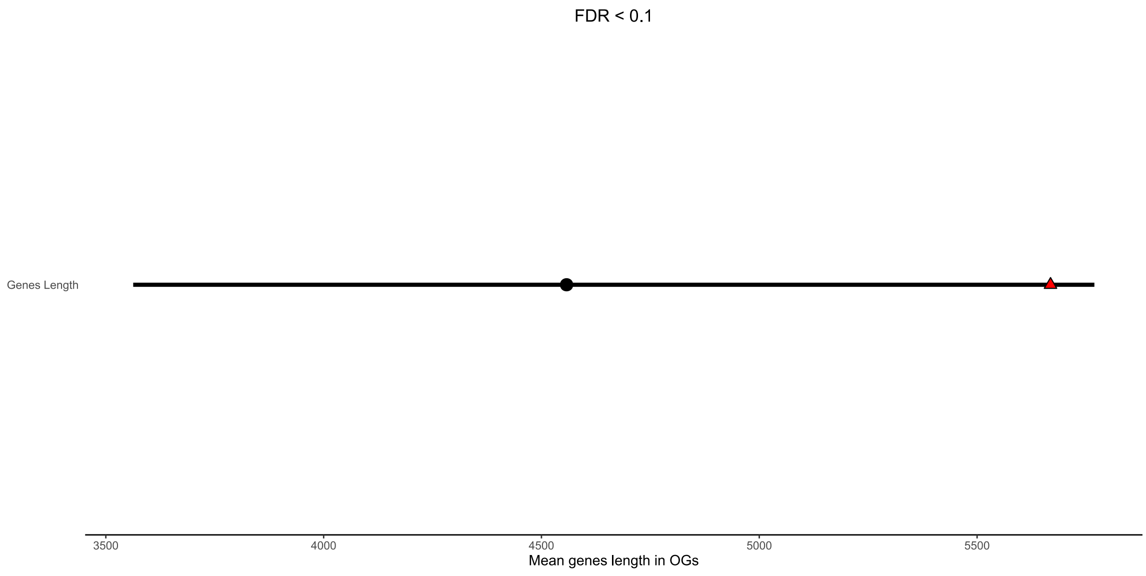


**Fig. S15.** Gene length bootstrapping assessment. The red triangle represents the mean gene length of the 33 candidates orthogroups (*PicMin*[1] FDR < 0.1). The black circle represents the mean of 10,000 random draws (of size 33). The black line represents the 95% confidence intervals of 10,000 random draws means.

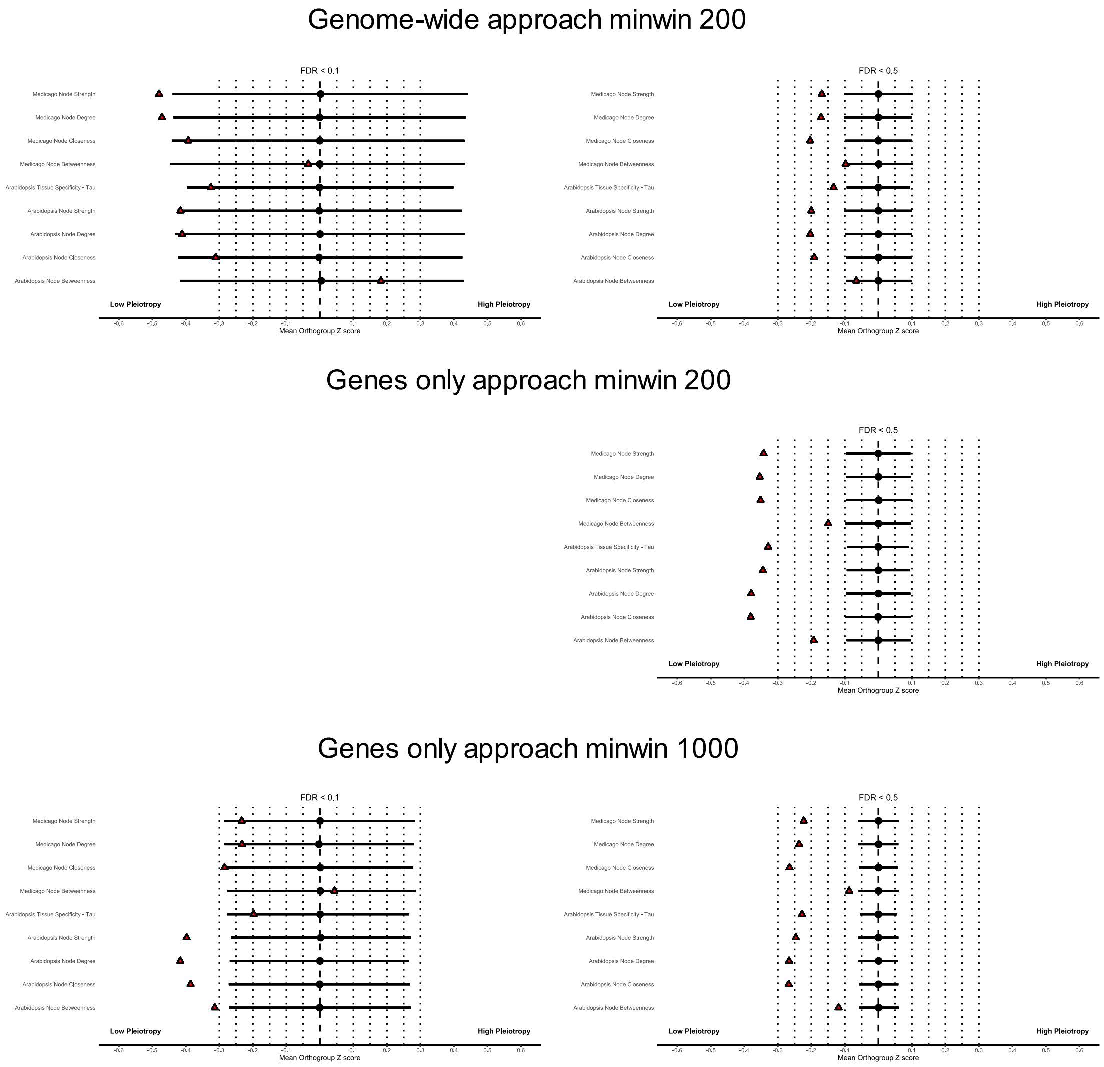


**Fig. S16.** Pleiotropy bootstrapping assessment for the *PicMin*[1] results derived from 3 alternative analysis approaches. Red triangles represent the average pleiotropy of the candidate orthogroups at the chosen *PicMin*[1] FDR thresholds, specified at the top of each plot (0.1,0.5). Black circles represent the average of 10,000 random draws means. Black lines represent the 95% interval of 10,000 random draws means. Rows in each plot correspond to the same assessment performed using different pleiotropy measures, labelled on the left. (FDR < 0.1 assessment not provided for the genes only minwin 200 approach due to the small size of the candidate orthogroups at that threshold).

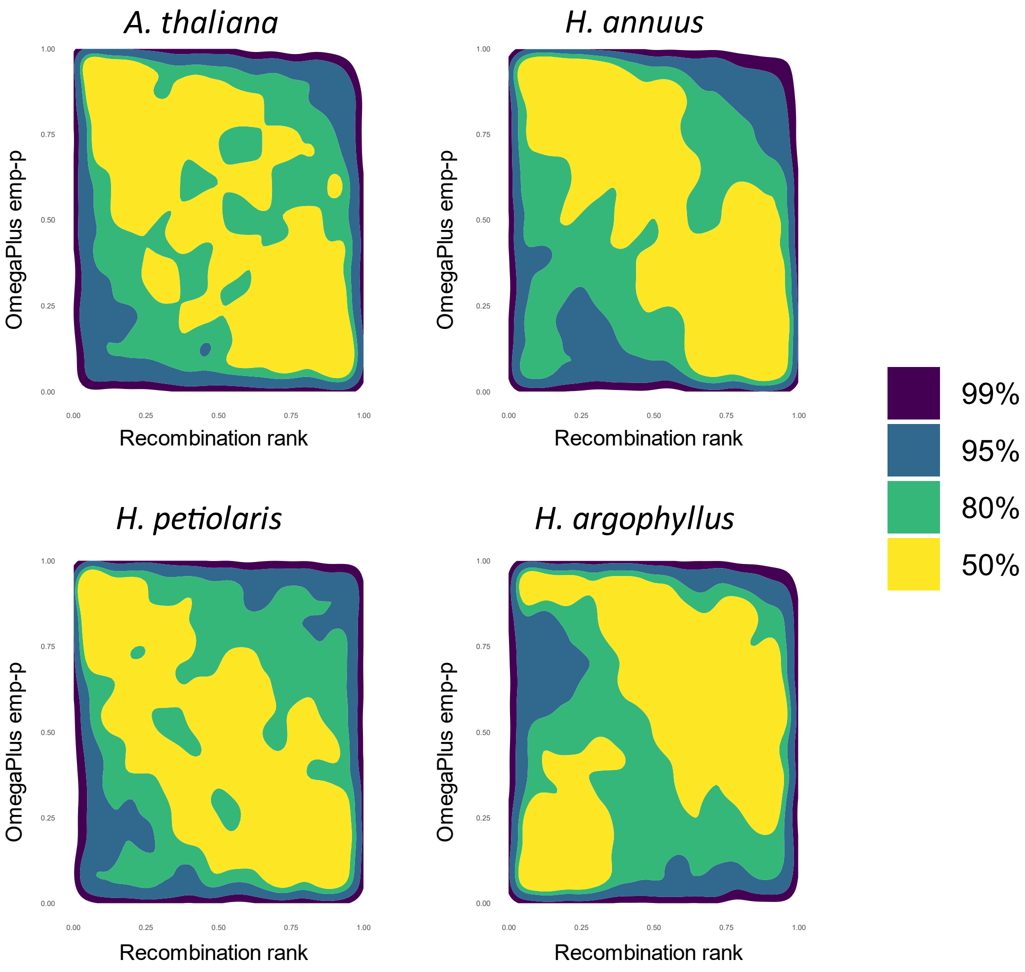


**Fig. S17.** Correlation between genes *OmegaPlus*[9] empirical p-values and genes recombination rate rank, with lower ranks corresponding to lower recombination rates. If no correlation exists, regions of high gene density (yellow) should be moderately even throughout the plotting area.

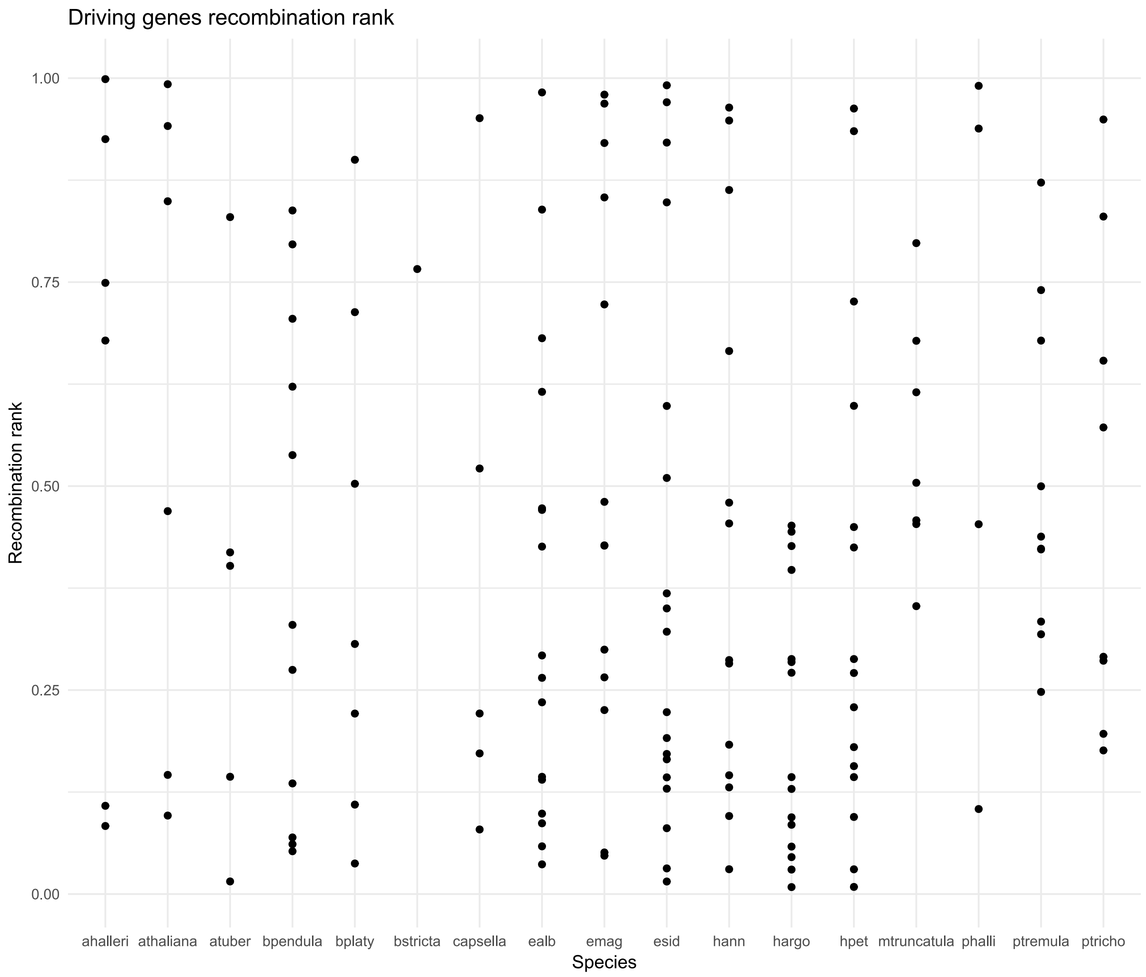


**Fig. S18.** Driving genes recombination rank within species. The plot displays the recombination ranks for the driving genes (*OmegaPlus*[9] empirical p-values < 0.1) of each species, for the 33 *PicMin*[1] orthogroups with FDR < 0.1. A lower recombination rank corresponds to a lower recombination rate.

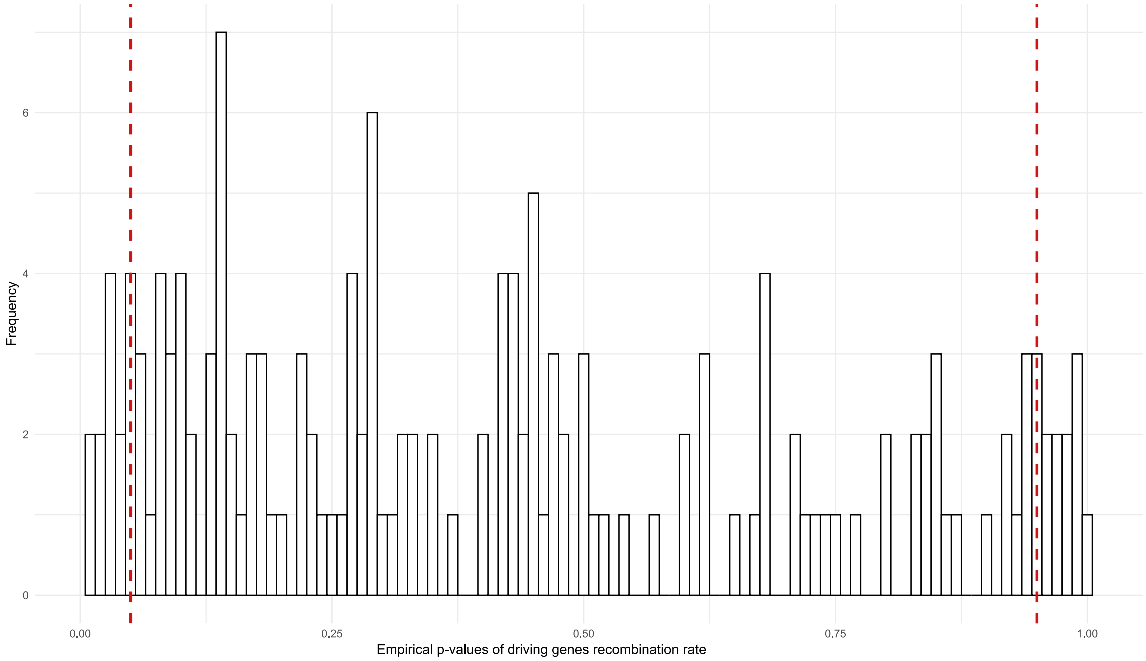


**Fig. S19.** Driving genes recombination rank distribution across species. The plot displays the distribution of the recombination ranks for the 160 driving genes (*OmegaPlus*[9] empirical p-values < 0.1 for the 33 *PicMin*[1] orthogroups with FDR < 0.1) for which we were able to calculate recombination rates. A lower recombination rank corresponds to a lower recombination rate. Red lines mark the 5% tails of the distribution: 0.05 and 0.95.

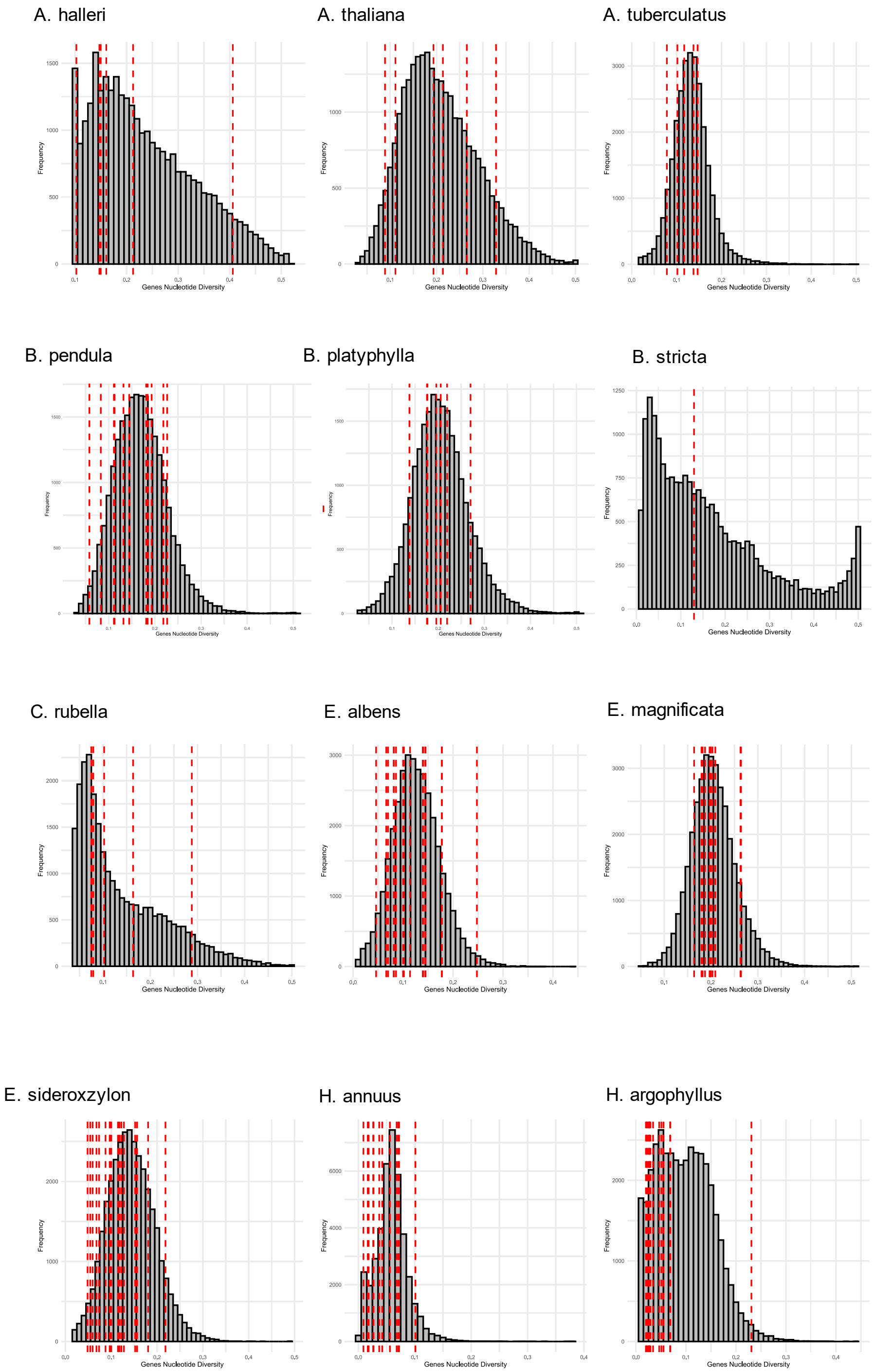
A. halleri
A. thaliana
A. tuberculatus
B. pendula
B. platyphylla
B. stricta
C. rubella
E. albens
E. magnificata
E. sideroxzylon
H. annuus
H. argophyllus
Frequency
Genes Nucleotide Diversity

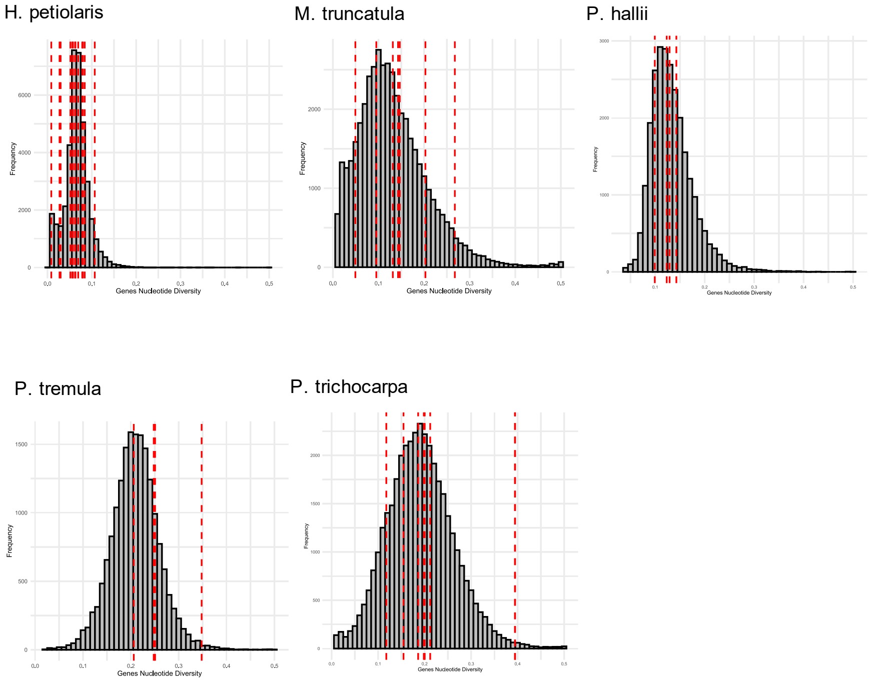


**Fig. S20.** Driving genes nucleotide diversity within species**.** Grey histograms plots showing the distribution of nucleotide diversity for all the genes in each species. Red lines mark the nucleotide diversity of driving genes (*OmegaPlus*[9] empirical p-values < 0.1 for the 33 *PicMin*[1] orthogroups with FDR < 0.1).

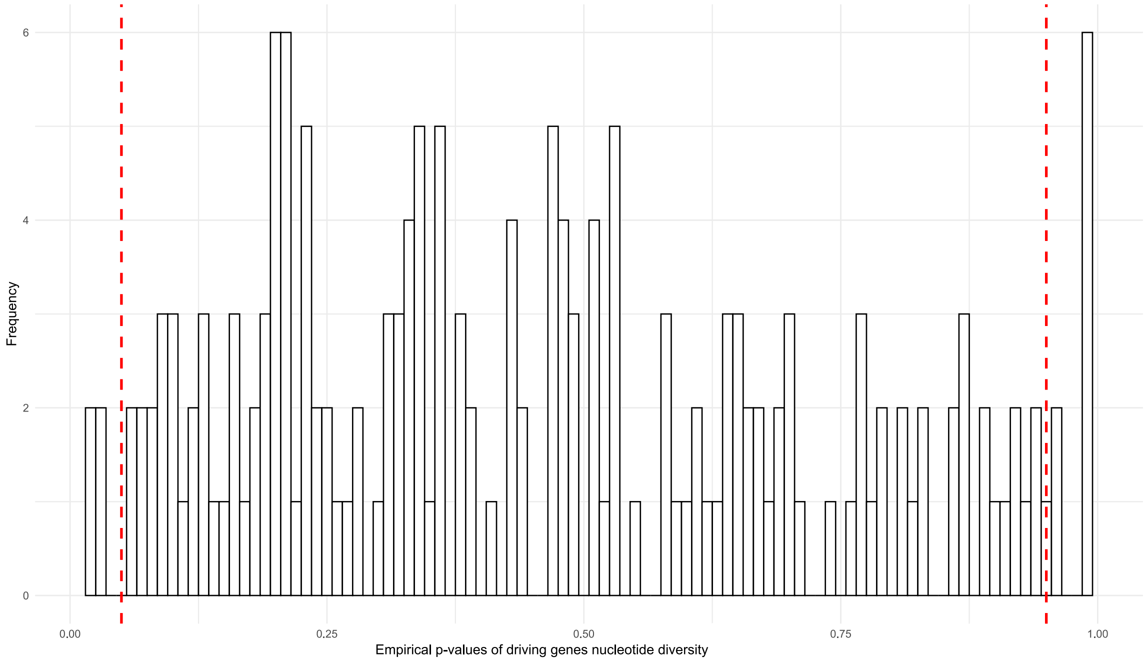


**Fig. S21.** Driving genes nucleotide diversity distribution across species. The plot displays the distribution of the nucleotide diversity empirical p-values for the 172 driving genes (*OmegaPlus*[9] empirical p-values < 0.1 for the 33 *PicMin*[1] orthogroups with FDR < 0.1). A lower empirical p-value corresponds to a lower nucleotide diversity. Red lines mark the 5% tails of the distribution: 0.05 and 0.95.

### Supporting Tables

**Table S1.** Summary table of the 17 datasets included in the analysis.

| Dataset | Reference genome resources (reference, GFF, proteome) | Individuals | Raw SNPs | Unrelated Individuals | Filtered SNPs | Ref. |
|---|---|---|---|---|---|---|
| Amaranthus tuberculatus | Amaranthus tuberculatus (Version: 2 54057) https://genomevolution.org/coge/SearchResults.pl?s=amaranthus%20tuberculatus&p=genome | 167 | 55,076,192 | 109 | 39,621,487 | 11 |
| Arabidopsis halleri | Arabidopsis halleri (Version: 2.2) https://www.ebi.ac.uk/ena/browser/view/GCA_900078215.1 | 55 | 7,736,676 | 20 | 2,485,101 | 12 |
| Arabidopsis thaliana | Arabidopsis thaliana (Version: TAIR10.1) https://www.ncbi.nlm.nih.gov/datasets/genome/GCF_000001735.4/ | 160 | 2,469,825 | 160 | 2,469,420 | 13 |
| Betula pendula | Betula pendula (Version: Bpev01) https://treegenesdb.org/org/Betula-pendula | 74 | 28,138,813 | 57 | 15,203,299 | 14 |
| Betula platyphylla | Betula pendula (Version: Bpev01) https://treegenesdb.org/org/Betula-pendula | 71 | 29,877,778 | 63 | 20,271,958 | 15 |
| Boechera stricta | Boechera stricta (Version: LTM_2.2) https://www.ncbi.nlm.nih.gov/datasets/genome/?taxon=72658 | 484 | 750,975 | 185 | 667,641 | 16 |
| Capsella rubella | Capsella rubella (Version: 1_0) https://www.ncbi.nlm.nih.gov/genome/annotation_euk/Capsella_rubella/100/ | 46 | 8,110,565 | 46 | 2,483,759 | 17 |
| Eucalyptus albens | Eucalyptus grandis (Version: 1_0) https://www.ncbi.nlm.nih.gov/genome/annotation_euk/Eucalyptus_grandis/100/ | 221 | 209,846,342 | 208 | 31,831,186 | 18 |
| Eucalyptus magnificata | Eucalyptus grandis (Version: 1_0) https://www.ncbi.nlm.nih.gov/genome/annotation_euk/Euc | 47 | 109,437,580 | 41 | 36,765,181 | 18 |

| | | | | | | |
|---|---|---|---|---|---|---|
| | alyptus_grandis/100/ | | | | | |
| Eucalyptus sideroxylon | Eucalyptus grandis (Version: 1_0) https://www.ncbi.nlm.nih.gov/genome/annotation_euk/Eucalyptus_grandis/100/ | 149 | 184,069,706 | 137 | 26,834,117 | 18 |
| Helianthus annuus | Helianthus annuus (Version: Ha412HOv2.0) https://sunflowergenome.org/assembly-data/ | 614 | 630,920,568 | 565 | 50,268,965 | 19 |
| Helianthus argophyllus | Helianthus annuus (Version: Ha412HOv2.0) https://sunflowergenome.org/assembly-data/ | 299 | 250,063,981 | 293 | 23,823,801 | 19 |
| Helianthus petiolaris | Helianthus annuus (Version: Ha412HOv2.0) https://sunflowergenome.org/assembly-data/ | 475 | 451,371,140 | 438 | 33,436,603 | 19 |
| Medicago truncatula | Medicago truncatula (Version: v4) https://www.ebi.ac.uk/ena/browser/view/GCA_000219495.2 | 174 | 53,346,118 | 166 | 5,787,032 | 20,21 |
| Panicum hallii | Panicum hallii (Version: 3.1) https://data.jgi.doe.gov/refine-download/phytozome?organism=Phallii&expanded=495 | 57 | 55,539,290 | 52 | 19,127,325 | 22 |
| Populus tremula | Populus tremula (Version: Potra02) https://www.biorxiv.org/content/10.1101/805614v1.full | 94 | 22,781,470 | 91 | 18,562,928 | 23 |
| Populus trichocarpa | Populus trichocarpa (Version: 210) https://phytozome-next.jgi.doe.gov/info/Ptrichocarpa_v3_0 | 191 | 17,232,173 | 182 | 13,846,730 | 24 |

**Table S2.** Summary table of annotation of the 15 high confidence orthogroups identified by intersecting the 33 candidate OGs identified in the main analysis (FDR < 0.1) with results derived from nine additional *PicMin* omitting closely related *Eucalyptus* and *Helianthus* species.

| **Orthogroup** | ***M. truncatula*/*A. thaliana* gene** | **Protein name** |
|---|---|---|
| OG0000525 | AT1G61680 | Terpene Synthase 14 |
| OG0001035 | AT4G24650 | Isopentenyltransferase 4 |
| OG0001601 | AT2G31960 | Callose Synthase 3 |
| OG0001991 | MTR_3g051230 | Cytochrome P450 |
| OG0002218 | AT5G14070 | ROXY1, ROXY2 |
| OG0002608 | AT1G11340 | G-type lectin S-receptor-like serine/threonine-kinase |
| OG0003537 | AT2G06925 | Phospholipase a2-alpha |
| OG0004282 | AT5G64620 | Vacuolar inhibitor of Fructosidase 2 |
| OG0004755 | AT1G16705 | p300/CBP acetyltransferase-related protein-like protein |
| OG0004877 | AT2G46940 | NA |
| OG0005418 | AT4G09170 | Transmembrane protein |
| OG0005857 | AT5G11000 | NA |
| OG0006228 | AT1G22810 | Ethylene response factor (ERF19) |
| OG0007151 | AT2G40640 | Plant U-box type E3 ubiquitin ligase (PUB62). |
| OG0010096 | AT2G20790 | Clathrin adaptor complexes medium subunit family protein |

**Table S3.** Correlations (*Pearson's r*) between number of duplications (based on our *Orthofinder2* assignment) and pleiotropy measures per orthogroup.

| **Pleiotropy measure** | **Pearson's r** | **p-value** |
|---|---|---|
| A. thaliana tissue specificity | 0.09 | < 2.2e-16 |
| *A. thaliana* betweenness | 0.06 | 1.637e-09 |
| *A. thaliana* strength | -0.19 | < 2.2e-16 |
| *A. thaliana* degree | -0.19 | < 2.2e-16 |
| *A. thaliana* closeness | -0.17 | < 2.2e-16 |
| *M. truncatula* betweenness | 0.10 | < 2.2e-16 |
| *M. truncatula* strength | -0.09 | < 2.2e-16 |
| *M. truncatula* degree | -0.10 | < 2.2e-16 |
| *M. truncatula* closeness | -0.09 | < 2.2e-16 |

**Table S4.** Correlations (*Pearson's r*) between the *OmegaPlus* empirical p-values obtained with the main approach (minimum window 500) and those obtained with three additional approaches (minimum window 200,1000 and genome-wide).

| **Species** | **200** | **1000** | **genome-wide** |
|---|---|---|---|
| *A. halleri* | 0.984377076862184 | 0.9432849 | 0.3458823 |
| *A. thaliana* | 0.938047193210014 | 0.8155351 | 0.383615 |
| *A. tuberculatus* | 0.809993464399262 | 0.6442576 | 0.3382122 |
| *B. pendula* | 0.791347812640607 | 0.5905464 | 0.3128254 |
| *B. platyphylla* | 0.81962796472042 | 0.6546213 | 0.3643874 |
| *B. stricta* | 0.998408462143016 | 0.9969038 | 0.1323291 |
| *C. rubella* | 0.93255478330479 | 0.7873099 | 0.4130388 |
| *E. albens* | 0.831974803206425 | 0.6908606 | 0.4172613 |
| *E. magnificata* | 0.786880772903479 | 0.6294256 | 0.3557499 |
| *E. sideroxylon* | 0.789363098364061 | 0.6024485 | 0.3698541 |
| *H. annuus* | 0.851460134833448 | 0.6783862 | 0.3462007 |
| *H. argophyllus* | 0.831802485515381 | 0.660224 | 0.3872186 |
| *H. petiolaris* | 0.867129116196599 | 0.7007766 | 0.3863179 |
| *M. truncatula* | 0.948138151401531 | 0.8091836 | 0.4352448 |
| *P. halli* | 0.70255852438136 | 0.4315459 | 0.194228 |
| *P. tremula* | 0.782778044371532 | 0.6136895 | 0.3515372 |
| *P. trichocarpa* | 0.869106003965276 | 0.6871917 | 0.3500445 |
